# A root-specific NLR network confers resistance to plant parasitic nematodes

**DOI:** 10.1101/2023.12.14.571630

**Authors:** Daniel Lüdke, Toshiyuki Sakai, Jiorgos Kourelis, AmirAli Toghani, Hiroaki Adachi, Andrés Posbeyikian, Raoul Frijters, Hsuan Pai, Adeline Harant, Karin Ernst, Martin Ganal, Adriaan Verhage, Chih-Hang Wu, Sophien Kamoun

## Abstract

Nucleotide-binding domain and leucine-rich repeat immune receptors (NLRs) confer disease resistance to a multitude of foliar and root parasites of plants. However, the extent to which NLR immunity is expressed differentially between plant organs is poorly known. Here, we show that a large cluster of tomato genes, which encodes the cyst and root-knot nematode disease resistance proteins Hero and MeR1 as well as the NLR-helper NRC6, exhibits nearly exclusive expression in the roots. This root-specific gene cluster emerged in *Solanum* species about 21 million years ago through gene duplication from the ancient NRC network of asterid plants. NLR-sensors in this gene cluster exclusively signal through NRC6 helpers to trigger the hypersensitive cell death immune response. These findings indicate that the NRC6 gene cluster has sub-functionalized from the larger NRC network to specialize for resistance against root pathogens, including cyst and root-knot nematodes. We propose that NLR gene clusters and networks have evolved organ-specific gene expression as an adaptation to particular parasites and to reduce the risk of autoimmunity.

## Introduction

Plants possess immune receptors that can detect invading pathogens and trigger potent immune responses. Intracellular nucleotide-binding domain and leucine-rich repeat (NLR) immune receptors are the most abundant class of plant resistance genes and confer immunity against various pathogenic microbes and parasitic pests, including nematodes and insects (Jones et al., 2016; Kourelis and van der Hoorn, 2018; Kourelis et al., 2021). NLRs recognize pathogen-secreted effector proteins and activate immune responses, typically culminating in a form of programmed cell death, known as hypersensitive response (Jones and Dangl, 2006). Pathogen effector recognition by NLRs can occur directly, through interactions between NLRs and cognate effector proteins, or indirectly, through guarding of host effector targets by NLRs (van der Hoorn and Kamoun, 2008). Despite significant advances in our understanding of NLR biology, fundamental questions about these receptors remain unanswered. For example, the degree to which NLR immunity is expressed differentially between plant organs is poorly understood (Deng et al., 2017; Barbey et al., 2019; Millett et al., 2015; Adachi et al., 2023). A meta-analysis of NLR gene expression patterns across plant species revealed distinct organ-specific expression profiles in monocot and Fabaceae plants, which primarily express NLR genes in roots, whereas Brassicaceae plants, such as Arabidopsis, show relatively higher NLR expression in shoots (Munch et al., 2018). Recently, a specific subset of NLR genes was shown to have a cell-specific, vasculature-enriched expression pattern, which is upregulated upon fungal infection (Tang et al., 2023). These findings suggest that plant cells in different organs, tissues and cell-types may have evolved to express distinct repertoires of NLR receptors as an adaptation to the parasites that attack specific host organs or cell-types (Adachi et al., 2019b; Munch et al., 2018; Tang et al., 2023). In this study, we address this question by demonstrating that an NLR gene network that confers resistance to plant-parasitic nematodes is specifically expressed in the roots of tomato.

While a number of NLRs function independently, encompassing both effector sensing and immune signaling in a single unit (functional singletons), many NLRs establish pairs or larger networks with sensor-helper relationships (Wu et al., 2017; Adachi et al., 2019b). One well-characterized example of networked NLRs are helper NRCs (NLRs required for cell death) and their disease resistance sensors, which form complex receptor networks of coiled-coil (CC) type NLRs (CC-NLRs) in the lamiid clade of asterid plants (Gabriëls et al., 2007; Wu et al., 2017; Kourelis et al., 2022; Sakai et al., 2023; Goh et al., 2023). Similar to paired NLRs, NRC sensor NLRs have specialized in the detection of pathogen effectors and require NRC helpers for immune responses and the induction of a hypersensitive cell death (Kourelis and Adachi, 2022; Contreras et al., 2023b). Upon pathogen activation, NRC sensors induce the oligomerization of NRC helpers through an activation-and-release model (Contreras et al., 2023a; Ahn et al., 2023). Activated NRCs subsequently assemble into higher order oligomers, similar to the wheel-like resistosome structures of singleton CC-NLRs, such as ZAR1 and Sr35 (Wang et al., 2019; Förderer et al., 2022; Ahn et al., 2023; Contreras et al., 2023a; Sakai et al., 2023).

ZAR1 and other CC-NLR resistosomes function as calcium ion (Ca^2+^) channels at the plasma membrane, an activity required for induction of the hypersensitive cell death (Wang et al., 2019; Jacob et al., 2021; Bi et al., 2021; Förderer et al., 2022). This activity depends on the very N-terminal α1 helix of the CC domain which assembles into a funnel-shaped structure that forms the resistosome membrane pore (Wang et al., 2019; Jacob et al., 2021; Bi et al., 2021; Adachi et al., 2019a; Förderer et al., 2022). The α1 helix is defined by the “MADA motif,” an ancient consensus sequence conserved in about 20% of CC-NLRs from dicot and monocot plants (Adachi et al., 2019a) and present also in bryophytes (denoted as MAEPL motif) (Chia et al., 2022). Remarkably, the MADA α1 helix motif has become non-functional in a large number of CC-NLRs, presumably because they have specialized into sensor NLR activities and rely on MADA containing helpers for induction of a hypersensitive cell death (Adachi et al., 2019a, 2019b). The NRC network adheres to this evolutionary “use-it-or-lose-it” model: Whereas the NRC helpers carry the consensus MADA sequence, the massively expanded phylogenetic NRC sensor clade lacks sequences matching this motif (Adachi et al., 2019a, 2019b; Contreras et al., 2023b; Sakai et al., 2023). In fact, a major subclade of NRC sensors carries the ∼400-1200 amino acid N-terminal Solanaceae Domain (SD) as an extension prior to the CC domain, a feature which would presumably preclude the formation of a ZAR1-type resistosome channel (Adachi et al., 2019c; Seong et al., 2020). The SD-CC-NLR sensor clade arose early in the NRC network evolution through integration of sequences of unknown origin (Seong et al., 2020). This clade includes well-studied disease resistance proteins such as Mi-1.2, Rpi-blb2, Sw5b, R1 and Prf (Mucyn et al., 2006; Milligan et al., 1998; Brommonschenkel et al., 2000; Ballvora et al., 2002; van der Vossen et al., 2005), as well as the cyst nematode resistance protein Hero (Ellis and Maxon Smith, 1971; Ernst et al., 2002) and the root-knot nematode resistance protein MeR1 (Frijters et al., 2021), both of which originate from the wild tomato species *Solanum pimpinellifolium*.

In plant genomes, NLR genes can be found as isolated genes (genetic singletons) or in gene clusters that vary from pairs to over a dozen genes (Michelmore and Meyers, 1998; Meyers et al., 2003). In Arabidopsis, around half of the NLR genes occur in gene clusters that primarily arose from tandem gene duplication or unequal crossing-over events (Meyers et al., 2003; Van de Weyer et al., 2019; Lee and Chae, 2020). Many pairs of sensor and helper NLRs are genetically clustered, often in head-to-head orientation, potentially to enable coordinated gene expression (Ashikawa et al., 2008; Narusaka et al., 2009; Okuyama et al., 2011; Césari et al., 2014; Huh et al., 2017; Białas et al., 2018). In contrast, functionally connected genes in the NRC network are not always genetically linked and can be dispersed across the genomes as in the case of tomato (Wu et al., 2017; Seong et al., 2020; Sakai et al., 2023).

Plant parasitic nematodes, including the potato cyst nematodes *Globodera rostochiensis* and *Globodera pallida* and the root-knot nematode *Meloidogyne enterolobii*, pose substantial threats to agriculture by inflicting root damage and reducing crop yields. Management of nematode diseases is particularly challenging due to their rapid reproduction and adaptability to various environments (Siddique et al., 2022). Their impact extends beyond agriculture, as they can harm natural ecosystems by disrupting plant root systems, altering ecosystem composition and functioning (Gillet et al., 2017; Li et al., 2023). Consequently, understanding the biology and ecology of plant-parasitic nematode interactions is essential for effective disease management strategies in agricultural and natural contexts.

Several nematode resistance genes belong to the SD-CC-NLR sub-clade within the wider NRC superclade. One example is tomato Mi-1.2, an economically important root-knot nematode resistance gene which is encoded within an NLR gene cluster (Milligan et al., 1998). *Mi-1.2*-mediated immune responses depend on the helper NLR NRC4, despite their lack of genetic linkage (Wu et al., 2017). In contrast, the cyst and root-knot nematode resistance genes *Hero* and *MeR1* reside in orthologous gene clusters of genetically linked NLRs in *S. pimpinellifolium* (Ellis and Maxon Smith, 1971; Ernst et al., 2002; Frijters et al., 2021). Given that Hero and MeR1 lack a MADA α1 helix sequence and carry an N-terminal SD domain, they are likely dependent on helper NLRs to function (Adachi et al., 2019a; Seong et al., 2020). However, the identity of their putative helper NLR(s) is yet to be determined, and it remains unknown whether the expression of these nematode resistance genes is root-specific.

In this study, we applied a computational pipeline to predict functional connections between NLR proteins based on the parameters of intergenic distance and phylogenomics. We hypothesized that mining genomes for gene clusters consisting of phylogenetically unrelated NLRs (mixed-clade gene clusters) would reveal previously unknown sensor-helper pairings. We applied this approach to tomato (*Solanum lycopersicum*), a species that carries ∼200 NLRs. Our analysis identified seven mixed-clade NLR gene clusters in tomato, one of which carries orthologs of the nematode resistance genes *Hero* and *MeR1*, along with ∼10 paralogous NLRs we called Hero-cluster NLRs (HCNs) and 1-2 genes of the NRC helper NLR clade we called NRC6. Remarkably, HCNs, including the *Hero* and *MeR1* resistance genes, exclusively depend on NRC6 helpers to trigger a hypersensitive cell death. Furthermore, RNA-sequencing (RNA-seq) data from root and leaf tissues revealed nearly exclusive baseline expression of *HCNs* and *NRC6* in tomato roots. These results suggest that the *HCN* and *NRC6* genes have evolved as a root-specific NRC sub-network that mediates nematode resistance in *Solanum* plants. Our findings offer novel insights into the evolution and function of NRC-network-mediated nematode resistance and underscore the emergence of organ-specific NLR immunity in plants.

## Results

### Three of seven tomato NLR mixed phylogenetic clade gene clusters include NRC helpers

We used a computational pipeline to predict sensor-helper NLR gene clustering based on two parameters: genetic distance and phylogenomics (Figure 1A; Sakai et al., 2023). Our hypothesis is that NLR gene clusters that consist of phylogenetically unrelated genes (mixed-clade clusters) include candidate sensor-helper relationships. To test this hypothesis, we analyzed the NLRome of tomato (*S. lycopersicum*). We used NLRtracker (Kourelis et al., 2021) to identify 197 NLRs from the genome annotations ITAG3.2 and NCBI RefSeq of tomato cv. Heinz 1706 (SL3.0) (Figure 1A; Supplementary Data 1) and assigned them to the seven major phylogenetic NLR groups based on prior classifications (Wu et al., 2017; Kourelis et al., 2021). The number of NLRs for these phylogenetic groups ranged from 3 (CC_R_ clade) to 11, 24, 29, 31, 49 and 53 (NRC-H, CC_G10_, NRC-S_Rx_, TIR, CC_other_, and NRC-S_SD_, respectively) (Figure 1B, C; Supplementary Data 1; Supplementary Dataset 1).

**Figure 1.**
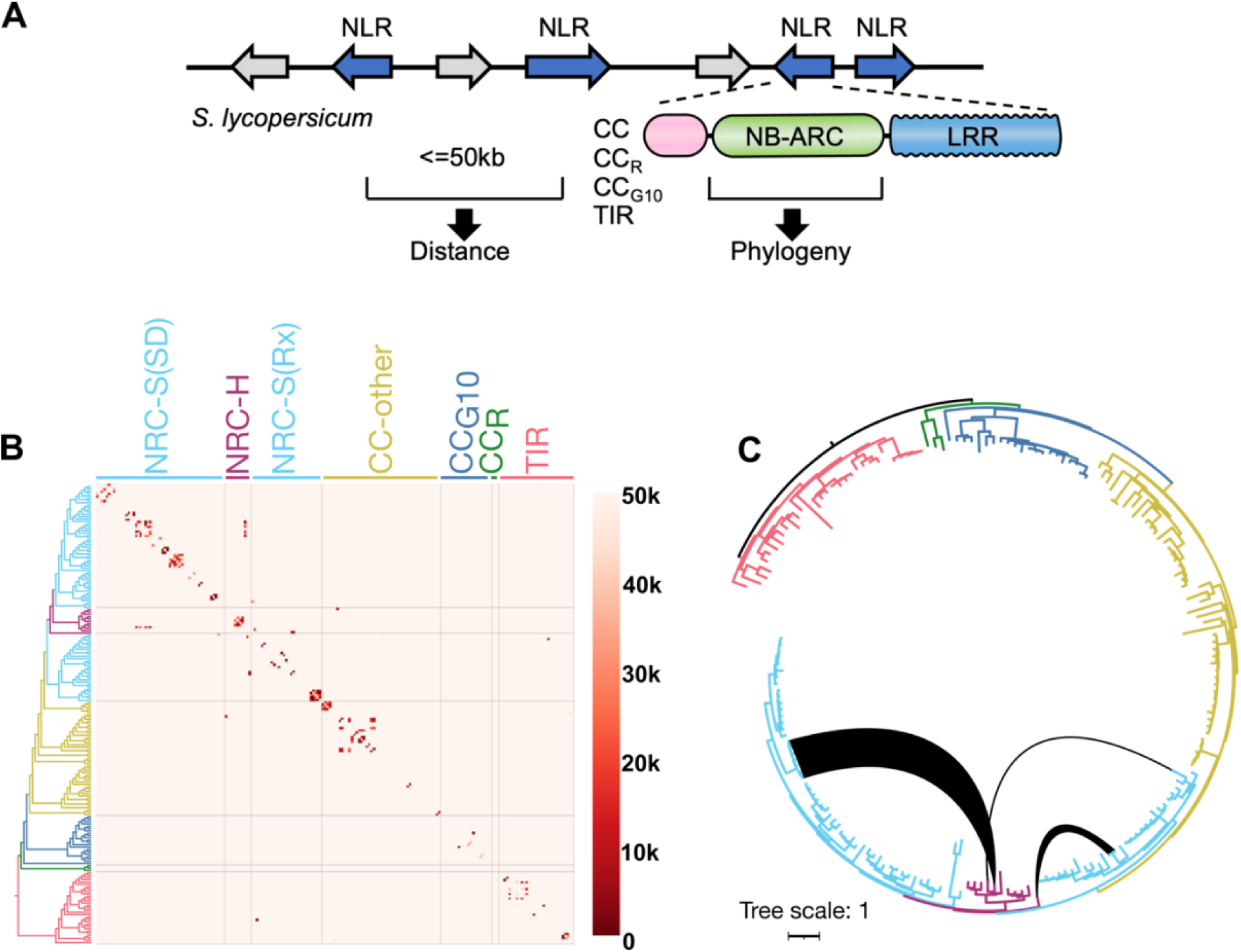
Tomato NLRs form clusters of phylogenetically related or unrelated NLR genes. **(A)** Schematic representation of the computational pipeline employed for predicting clustering of sensor-helper NLRs. The genetic distances among tomato (*S. lycopersicum*) NLR genes and their phylogenetic relationships were assessed using a custom Python script available at https://github.com/slt666666/gene-cluster-matrix. **(B)** NLR matrix integrating gene distance and phylogenetic relationships. Genetically linked NLR genes (distance < 50kb) are highlighted in the matrix, with colors representing gene distance and ordering based on phylogeny. The branches of major phylogenetic NLR clades (Wu et al., 2017; Kourelis et al., 2021) are color-coded, and NRC helpers (NRC-H) and NRC sensors (NRC-S) are depicted in red and blue, respectively. The NRC sensor clade is further subdivided into Rx-type (Rx) or Solanaceous domain (SD)-containing NRC sensors. An interactive version of the matrix is provided as Supplementary Data 2. **(C)** Phylogenetic tree with black lines connecting NRC helper and NRC sensor NLRs encoded in gene clusters containing phylogenetically unrelated NLRs (mixed-clade gene clusters). Gene distance information, alignments, and phylogenetic tree files are available as Supplementary Dataset 1.

To determine the genomic distribution of the 197 tomato NLRs, we calculated the distance between each of the NLR-encoding genes (Figure 1A, Supplementary Dataset 1) and created a genetic distance matrix in which the NLRs were grouped by phylogeny (Figure 1A, B; Supplementary Data 2; 3). Of the 197 NLRs, 107 (54%) were within 50 kb of another NLR gene and deemed to occur in 37 gene clusters consisting of 2 to 7 NLR genes per cluster (Figure 1B; Supplementary Figure 1A; Supplementary Data 3).

Next, we determined which NLR gene clusters contain mixed-clade genes. As evident from the genetic distance matrix, the great majority of NLR genes clustered with phylogenetically related NLRs, indicating the prevalence of tandem duplications during NLR gene expansion (Figure 1B; Supplementary Data 2; 3). A total of 30 out of 37 NLR gene clusters included NLRs that belong to the same phylogenetic clade (Supplementary Figure 1A; Supplementary Data 2; 3). However, we identified seven mixed-clade NLR gene clusters (Figure 1; Supplementary Figure 1, Supplementary Data 2; 3). Notably, three of these gene clusters contain NRC helpers and NLRs from the NRC-dependent sensor clades, previously defined as NRC-S_Rx_ and NRC-S_SD_ (Figure 1B, C; Supplementary Figure 1; Supplementary Data 2; 3; Wu et al., 2017; Kourelis et al., 2021). We considered NLRs encoded in these mixed-clade gene clusters as potential candidates for sensor-helper pairs or networks of the NRC phylogenetic clade.

### The cyst nematode resistance protein Hero is encoded in a large NRC mixed-clade gene cluster present across *Solanum* species

We noted that the largest mixed-clade NLR gene cluster identified contains homologs of the previously identified resistance gene *Hero* that functions against the potato cyst nematodes *G. rostochiensis* and *G. pallida* (Ellis and Maxon Smith, 1971; Ernst et al., 2002), and an uncharacterized NRC helper clade protein (Figure 1B, C; Supplementary Figure 1; Supplementary Data 2; 3). Hero belongs to the branch of the phylogenetic NRC superclade that includes NLRs with the Solanaceae Domain (SD) N-terminal extension prior to the CC domain (NRC-S(SD) in Figure 1B; Supplementary Data 2; 3; Supplementary Dataset 1). We reasoned that this gene cluster may consist of sensor-helper NLRs and further investigated this hypothesis.

The NRC helper clade protein encoded in this gene cluster is one of 11 NRC helper superclade proteins in tomato and was previously defined as NRC6 (Solyc04g008150.2; Wu et al., 2017). To determine the phylogenetic relationship of NRC6 and the genetically clustered Hero homologs, we constructed a phylogenetic tree with 20,292 NLR sequences extracted from a database derived from 124 plant genomes (Figure 2A, top left; Supplementary Data 4). NRC6 belongs to a phylogenetic clade that includes the uncharacterized NRC family proteins NRC5, NRC7 and NRC8 and is more closely related to the functionally characterized NRC4 sub-clade than to the NRC1, NRC2, NRC3 and NRCX sub-clade (Figure 2A, bottom left; Wu et al., 2017; Kourelis et al., 2021). Tomato NRC6 and the closely related NRC7 contain an atypical 12 amino acid N-terminal extension before their MADA motif, which scores lower compared to motifs of other functionally characterized NRC proteins when applying a Hidden-Markov Model for MADA motif detection (Supplementary Figure 2; Adachi et al., 2019a). However, NRC6 is predicted to be a canonical helper NLR protein, containing an intact P-loop and MHD motif (Supplementary Figure 2).

**Figure 2.**
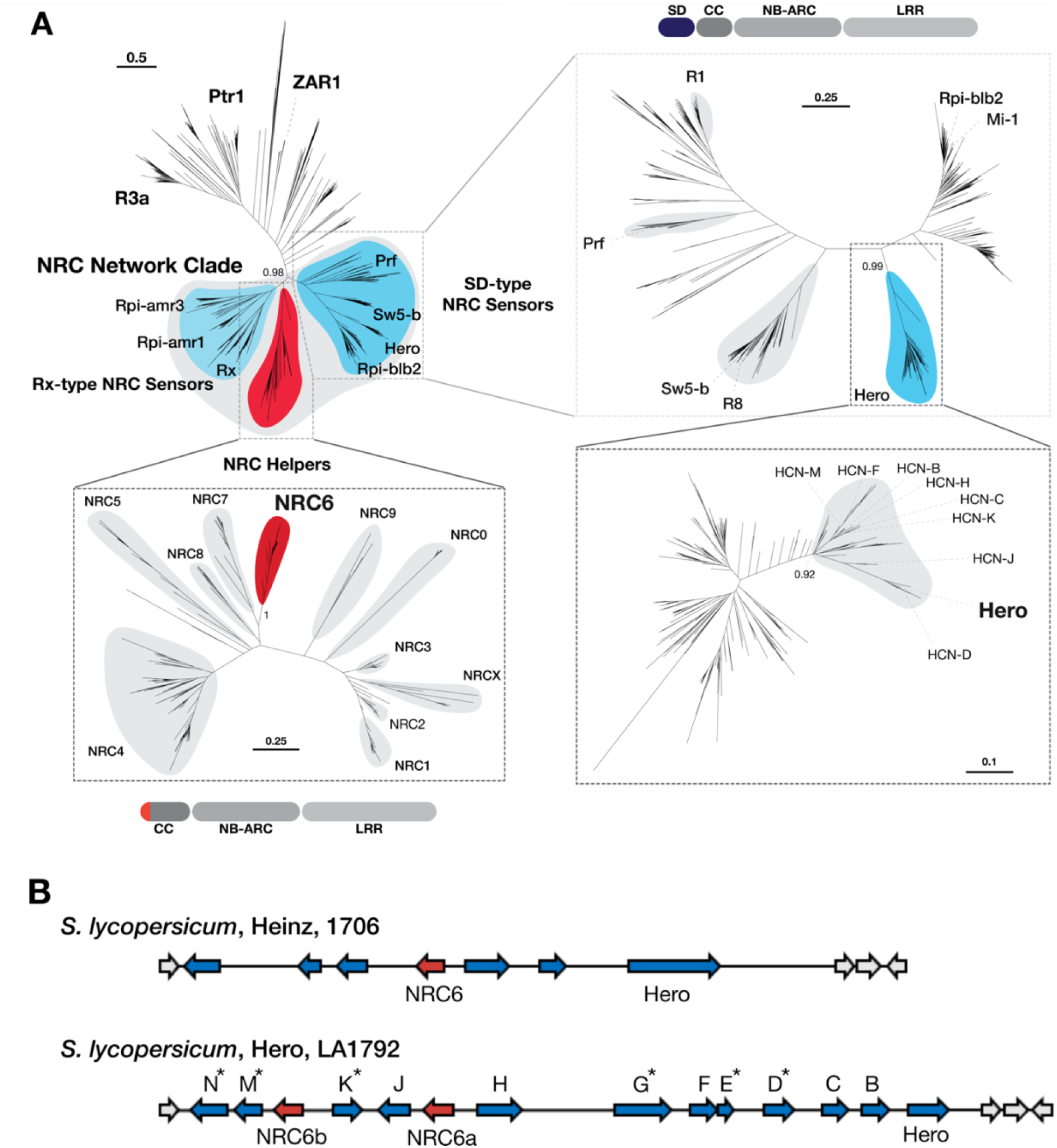
Hero and NRC6 form distinct phylogenetic subclades within the NRC helper and SD-type NRC sensor clades. **(A)** The phylogenetic tree of NLRs was constructed based on the NB-ARC domain from a large NLR sequence database (Supplementary Dataset 2). A total of 20.292 NLR sequences was aligned using MAFFT and the phylogeny was inferred using FastTree to determine the NRC network clades of NRC helpers, Rx-type NRC sensors and SD-type NRC sensors **(top left)**. Subtrees for the NRC helper **(bottom left)** and SD-type NRC sensor **(top right)** subclades were generated, containing well-defined phylogenetic clades including the Hero resistance protein and NRC6, respectively. The Hero-cluster NLR (HCN) clade is further divided into a tomato subclade (gray) which contains branches defined by proteins encoded in the NRC6 and HCN containing mixed-clade gene cluster **(bottom right)**. **(B)** Schematic depiction of the NRC6 and HCN encoding mixed-clade gene clusters of *S. lycopersicum* Heinz 1706 and Hero LA1792, which contains an introgressed cluster from *S. pimpinellifolium* LA121 (Ellis and Maxon Smith, 1971; Ernst et al., 2002). The HCNs encoded in this gene cluster define the branches of the tomato subclade shown in **(A, bottom right)**. Pseudogenes are indicated by an asterisk. Sequences, alignments, and phylogenetic tree file are available as Supplementary Dataset 2 or https://github.com/amiralito/Hero.

In the phylogenetic trees, the Hero resistance protein and other NLRs encoded in the *Hero* encoding gene cluster form a distinct phylogenetic clade (Hero-cluster NLR (HCN) clade) that is closely related to the phylogenetic clade containing the nematode resistance protein Mi-1.2 (Milligan et al., 1998) and the potato late blight resistance protein Rpi-blb2 (van der Vossen et al., 2005) (Figure 2A, top right). Mi-1.2, Rpi-blb2, and the HCNs all contain an N-terminal Solanaceae domain (SD) prior to the CC domain. Strikingly, the NRC6/HCN gene cluster is only detected in genomes of *Solanum* species, but is absent in the genomes of other Solanaceae, such as pepper (*Capsicum* spp.) or tobacco (*Nicotiana* spp.) (Supplementary Figure 3, Supplementary Data 4). We can therefore date the emergence of the NRC6/HCN gene cluster to around 17-21 MYA (Supplementary Figure 3) based on divergence date estimates (Särkinen et al., 2013).

In the Heinz reference tomato genome assembly with the ITAG3.2 annotation, the NRC6/HCN gene cluster is annotated to contain six potential HCN genes of the phylogenetic NRC sensor clade and one NRC helper clade encoding gene, arranged in two blocks of tandemly repeated NLRs (Figure 2). The *Hero* resistance gene was identified in an introgression line of cultivated tomato (*S*. *lycopersicum*, LA1792) and in the wild tomato species *S. pimpinellifolium* LA121 (Ellis and Maxon Smith, 1971; Ernst et al., 2002). The introgressed wild tomato NLR gene cluster encodes 12 HCNs that are phylogenetically related to Hero, some of which appear to be pseudogenes (Figure 2; Ernst et al., 2002). In addition, the gene cluster encodes for two NRC helper proteins, which phylogenetically cluster with NRC6 and were therefore termed NRC6a and NRC6b (Figure 2). The comparison between the Heinz *S*. *lycopersicum* and the Hero *S. pimpinellifolium* NRC6/HCN gene cluster provides a glimpse of this cluster’s sequence variation between *Solanum* species (Figure 2B).

We compared the NRC6/HCN gene cluster extracted from chromosome-scale genome sequence assemblies of the *Solanum* species wild tomato (*S. pennelli*), potato (*S. tuberosum*), and American black nightshade (*S. americanum*), to the *S*. *lycopersicum* and the *S. pimpinellifolium* gene clusters. This revealed significant genomic rearrangements, presence/absence polymorphisms and duplications of the encoded NLR genes (Supplementary Figure 4). In general, the extracted NRC6 sequences form a well-defined phylogenetic clade, whereas the HCNs appear more divergent, splitting into multiple subclades, as previously reported for NRC sensors of the SD clade (Figure 2A, bottom; (Seong et al., 2020). However, the conservation of the NRC6/HCN gene cluster across multiple *Solanum* species (Supplementary Figure 3; 4) suggests that the gene cluster encoded NRC6 and HCN proteins could be engaged in sensor-helper functional connections required for immune responses.

### Hero-cluster NLRs (HCNs) require NRC6 helper NLRs to induce a hypersensitive cell death

We hypothesized that Hero and other HCNs require their genetically clustered NRC6 for the induction of a hypersensitive cell death. To experimentally test this hypothesis, we used a genetic complementation assay based on transient expression by agroinfiltration in *Nicotiana benthamiana*, a species which lacks a functional NRC6 ortholog (Figure 3; Supplementary Figure 3; Adachi et al., 2023). We first cloned Hero and five of its HCN paralogs along with NRC6a or NRC6b, extracted from the *S. pimpinellifolium* Hero gene cluster sequence of the introgression tomato line LA1792 (Ellis and Maxon Smith, 1971; Ernst et al., 2002). We introduced a histidine (H) to alanine (A) mutation in the MHD motif of Hero and the other HCNs to generate autoactive mutants (van Ooijen et al., 2008) of the NLRs (referred to as MAD mutants or HCN^HA^). In the co-expression assays, Hero caused macroscopic cell death in *N. benthamiana* leaves when co-expressed as MAD mutant with NRC6b, but not with NRC6a (Figure 3B; Supplementary Figure 5; 6). In contrast, four of the five HCNs: HCN-B, HCN-F, HCN-H and HCN-J, caused cell death when co-expressed as MAD mutants with either NRC6a or NRC6b (Figure 3B; Supplementary Figure 5; 6). Among these HCNs, HCN-B also triggered cell death when co-expressed with NRC6a or NRC6b as both wildtype and MAD mutant variant (Figure 3B). Of the five tested HCNs, HCN-C did not induce cell death either as wildtype or autoactive MAD mutant when co-expressed with NRC6a or NRC6b (Figure 3B). However, in all cases, neither Hero nor the four functional HCNs caused cell death in the absence of the NRC6 proteins, indicating that these sensor-like autoactivated NLRs cannot signal for cell death without their helper NRC6 in *N. benthamiana* (Figure 3; Supplementary Figure 5). We conclude that Hero and the HCNs are functionally connected with their genetically linked NRC6a and NRC6b helper NLRs and form a new branch of the NRC network.

**Figure 3.**
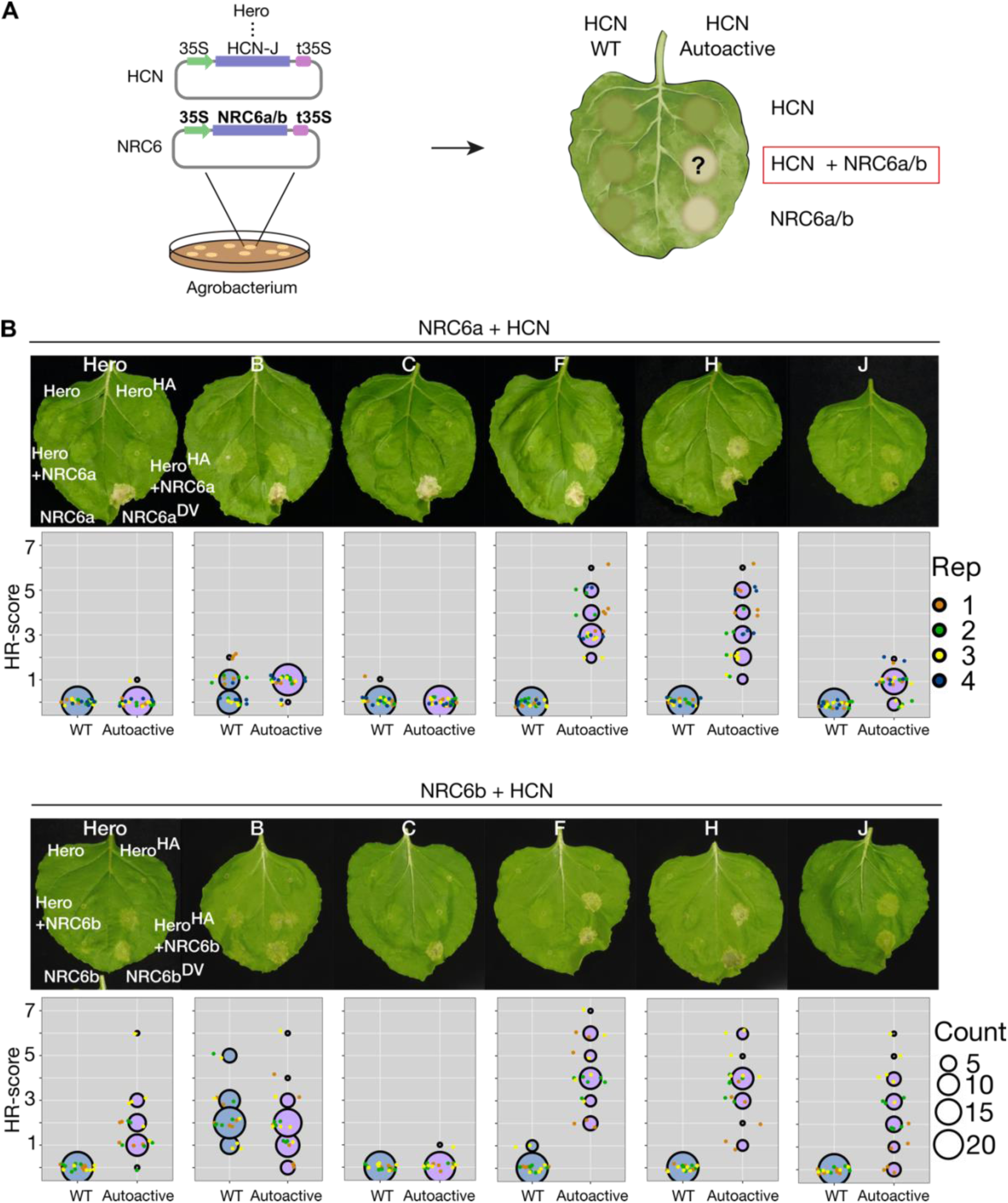
Hero-cluster NLRs (HCNs) require NRC6 helper NLRs to trigger a hypersensitive cell death in *Nicotiana benthamiana*. (A) The schematic representation illustrates the genetic complementation scheme employed throughout this experiment. Agrobacteria carrying NRC6a, NRC6b, or the specified HCN expression constructs were infiltrated into *N. benthamiana* leaves to express wildtype or autoactive HCNs (autoactive = HA) and NRC6 helper NRCs (autoactive = DV) in the indicated combinations. **(B)** Representative cell death phenotypes in *N. benthamiana* leaves induced by HCNs when co-expressed with NRC6a or NRC6b, photographed at 7 days post-infiltration (dpi) with agrobacteria. A p19 silencing construct was co-expressed in every infiltration. Cell death was visually scored and statistically analyzed. Data points are depicted as dots, with each biological replicate represented by a different color. The central circle for each cell death category proportionally represents the total number of data points for each treatment. Scoring is shown only for wildtype and autoactive HCNs, co-expressed with wildtype NRC6a or NRC6b helpers. Complete quantification and statistical analysis are presented in Supplementary Figure 5.

### NRC2, NRC3 and/or NRC4-dependent disease resistance proteins do not signal through NRC6

Our finding that the HCN and NRC6 gene cluster forms an NLR receptor sub-network within the much larger phylogenetic NRC superclade prompted us to determine the degree to which the network branches are functionally connected. To investigate this, we co-expressed with NRC6b the disease resistance proteins R1, Rpi-blb2, Mi-1.2, CNL11990, Pto, Gpa2, R8, Rx and Sw5b, all of which are known to be functionally dependent on NRC2, NRC3 and/or NRC4 (Wu et al., 2017). These NRC2/3/4- dependent sensors were either activated with their respective AVR effectors or expressed as autoactive mutants. In these assays, none of the nine NRC2/3/4-dependent NLR proteins caused a visible cell death response in the presence of NRC6b, unlike the cell death response they triggered in the presence of the positive controls NRC3 or NRC4a (Figure 4; Supplementary Figure 7). We conclude that NRC6 forms a helper node in the NRC superclade network that is distinct from the previously characterized NRC2, NRC3 and NRC4 nodes. In addition, we also conclude that resistance to parasitic nematodes in the NRC network is mediated by multiple NRC helpers, given that, unlike Hero, the NRC2- and NRC3-dependent potato cyst nematode resistance protein Gpa2 and the NRC4-dependent root-knot nematode resistance protein Mi-1.2 function independently of NRC6 (Figure 4).

**Figure 4.**
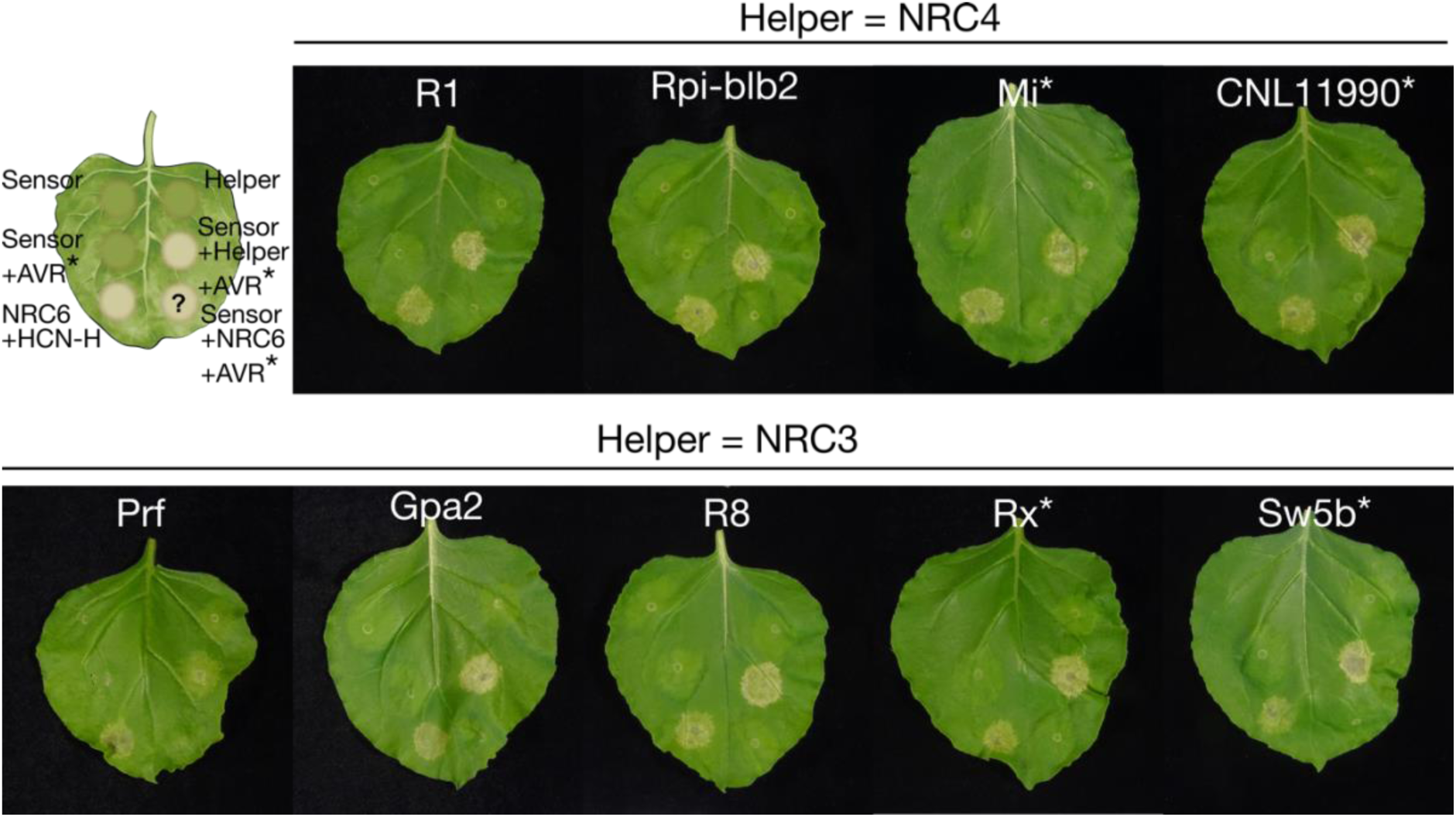
NRC2, NRC3 and/or NRC4-dependent disease resistance proteins do not signal through NRC6. The schematic representation illustrates the genetic complementation scheme employed throughout this experiment. Agrobacteria carrying expression constructs for sensors, helpers, or AVR effector genes were infiltrated in indicated combinations into the leaves of *N. benthamiana nrc2/3/4* mutant plants to trigger cell death. For Mi-1.2, CNL11990, Rx, and Sw5b, autoactive sensor mutants were expressed instead of a corresponding AVR effector gene (indicated by an asterisk). Tomato NRC3 or NRC4a was co-expressed as a positive control for complementation, respectively (Wu et al., 2017). HCN-H was used as a positive control for NRC6b-dependent cell death. Leaf images were photographed at 7 days post-infiltration (dpi) with Agrobacteria, a p19 gene silencing construct was co-expressed in each infiltration. Quantification and statistical analysis are presented in Supplementary Figure 7.

### Hero and the HCNs do not signal through the other nine tomato NRC helpers

Apart from NRC6a, NRC6b, and the NLR modulator NRCX (Adachi et al., 2023), there are nine additional NRC helper proteins in the tomato NRC phylogenetic clade (Supplementary Figure 2). To elucidate the relationship between NRC6 and other NRC clade proteins and to define the architecture of the NRC network, we co-expressed Hero and its five HCN paralogs with the other nine tomato NRCs (Figure 5). This enabled us to determine whether any of the HCN proteins could activate any of the other tomato NRC helpers. We co-expressed, Hero and each autoactive HCN with each tomato NRC helper, and used NRC6b as a positive control. We found that neither Hero, nor the other HCNs, could induce cell death through any other NRC proteins, except NRC6b (Figure 5; Supplementary Figure 8; 9). As an additional positive control, we co-expressed the NRC sensor Rx with each of the tomato NRC helpers, confirming that Rx can activate NRC2, NRC3, variants of NRC4, as well as NRC1 from tomato, consistent with previous reports (Figure 5; Supplementary Figure 8; Gabriëls et al., 2007; Wu et al., 2017)). It should be noted that although the tested NRC proteins expressed well in these experiments, we did not detect the accumulation of the NRC5 protein (Supplementary Figure 9). In summary, Hero, HCNs and NRC6 form a specific NRC sub-network that is phylogenetically related, but distinct from the major NRC2, NRC3 and NRC4 network.

**Figure 5.**
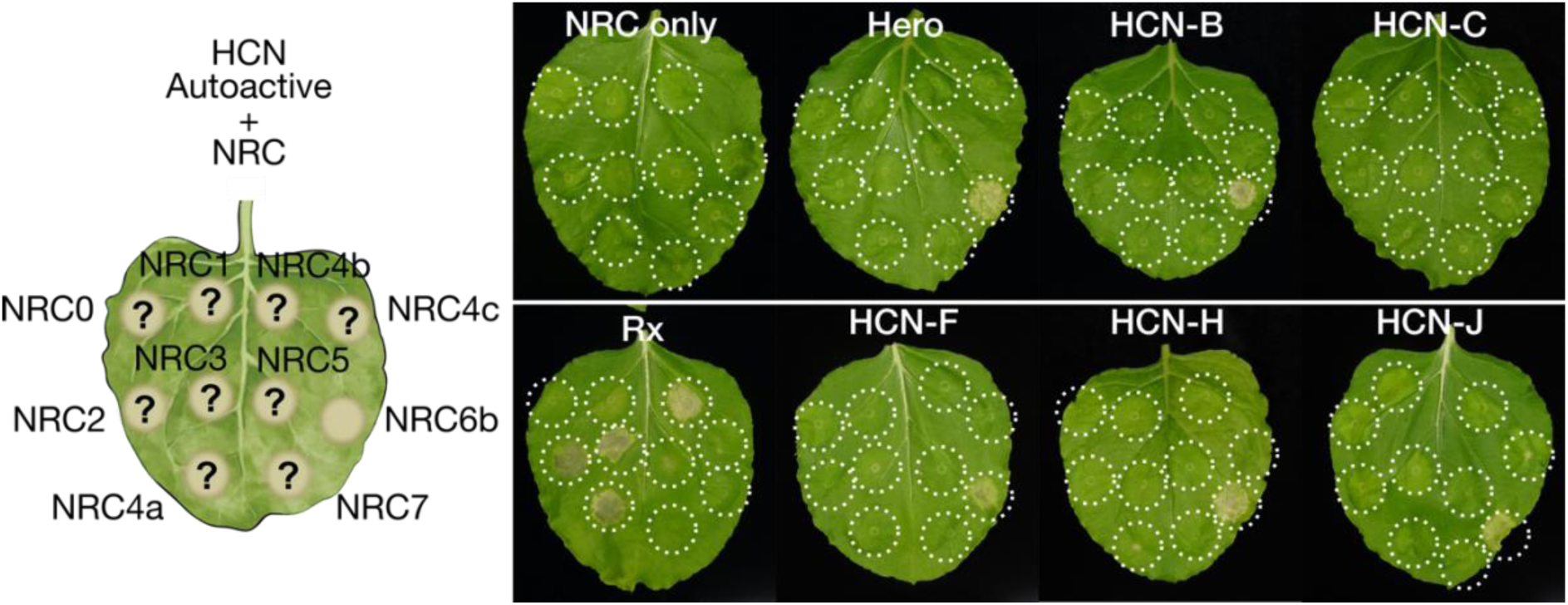
Hero and the HCNs do not signal through the other nine tomato NRC helpers. The schematic representation illustrates the genetic complementation scheme employed throughout this experiment. Agrobacteria carrying autoactive HCNs and the indicated wildtype tomato NRC helper expression constructs were co-infiltrated into the leaves of *N. benthamiana nrc2/3/4* mutant plants. As a positive control, Rx was co-expressed with all tomato NRC helpers. Leaves were photographed at 7 days post-infiltration (dpi) with Agrobacteria, a p19 silencing construct was co-expressed for each infiltration. Quantification and statistical analysis are presented in Supplementary Figure 8.

### The root-knot nematode resistance gene *MeR1* is a Hero-cluster NLR (HCN) that signals through NRC6

In addition to the cyst nematode resistance protein Hero (Ellis and Maxon Smith, 1971; Ernst et al., 2002), MeR1, a resistance protein against the root-knot nematode *M. enterolobii*, is also encoded within an orthologous HCN and NRC6 containing gene cluster in the wild tomato *S. pimpinellifolium* (Frijters et al., 2021). Based on phylogenetic analyses of HCN proteins, MeR1 is phylogenetically closest to HCN-F, raising the hypothesis of its specific dependence on NRC6 for cell death induction (Figure 6A). To examine this hypothesis, we co-expressed MeR1 in *N. benthamiana* leaves, both in its wildtype and autoactive histidine to alanine (MHD to MAD, Mer1^HA^) forms, with NRC6b or other tomato NRC helpers. Our results revealed that, similar to other HCNs, MeR1 exclusively signals through NRC6b, exhibiting no functional interaction with any of the other tomato NRC helpers we tested (Figure 6B; Supplementary Figure 10). In contrast, wildtype MeR1 or the autoactive MAD mutant did not induce cell death in *N. benthamiana* when expressed on their own (Figure 6B; Supplementary Figure 10). In these experiments, wildtype MeR1 displayed autoactivity when co-expressed with NRC6b, independently of the MAD mutation (Figure 6B; Supplementary Figure 10). We conclude that the NRC6/HCN gene cluster encodes resistance genes that function against two species of plant parasitic nematodes (Ernst et al., 2002; Frijters et al., 2021).

**Figure 6.**
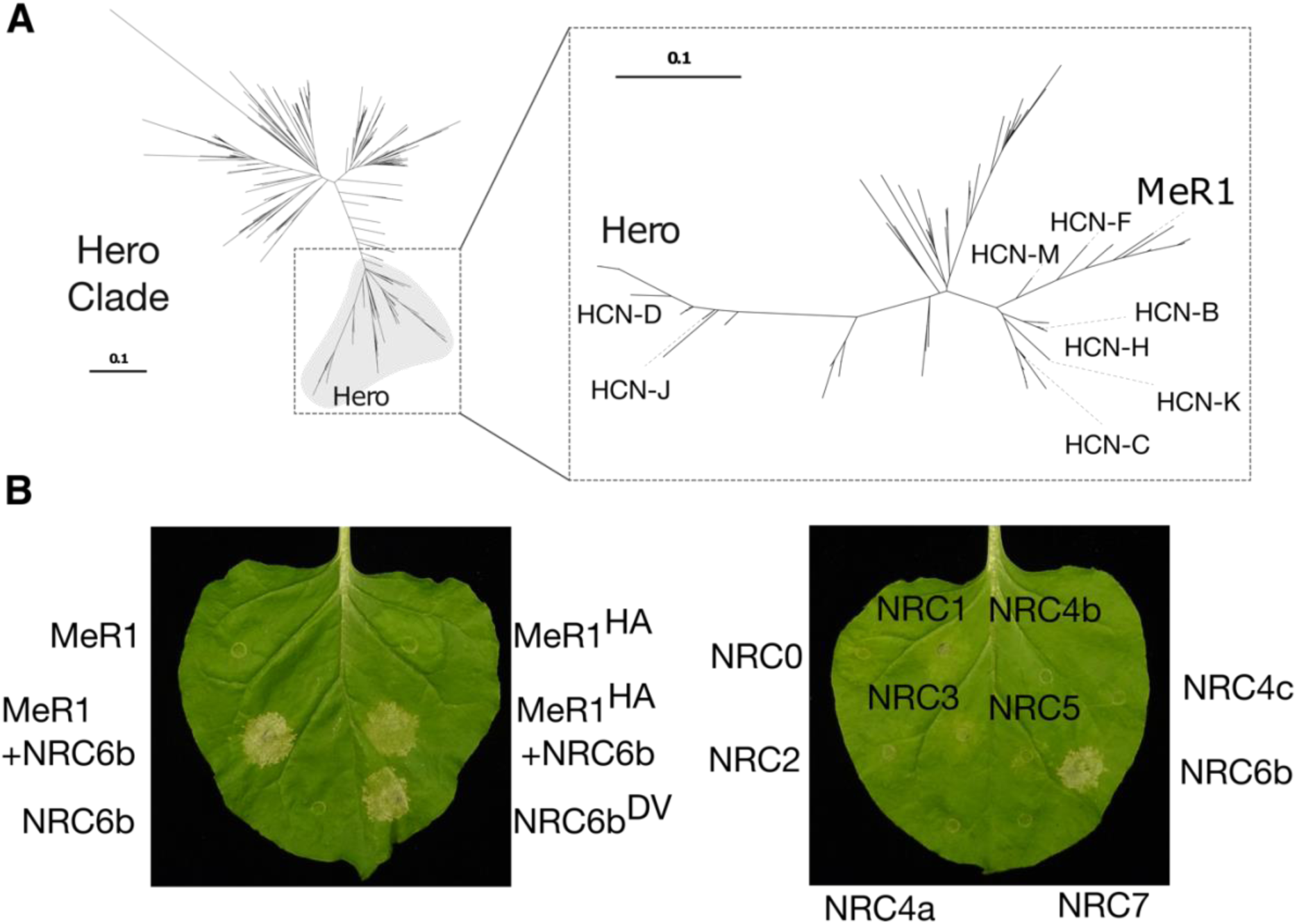
The root-knot nematode resistance gene MeR1 is a Hero-cluster NLR (HCN) that requires NRC6 to induce a hypersensitive cell death. **(A)** A phylogenetic sub-tree was generated with MeR1 and tomato and potato NLRs defined as Hero-cluster NLRs (HCNs), from data presented in Figure 2A. Re-alignment was performed using MAFFT, the phylogeny was inferred using FastTree. **(B)** Agrobacteria harboring wildtype or autoactive MeR1, or NRC helpers, were infiltrated in the indicated combinations into leaves of *N. benthamiana* wildtype (left panel) or *nrc2/3/4* mutant (right panel) plants for co-expression. Leaves were photographed at 7 days post-infiltration (dpi) with Agrobacteria, a p19 silencing construct was co-expressed for each infiltration. Quantification and statistical analyses are presented in Supplementary Figure 10. Sequences, alignments and the phylogenetic tree file are available as Supplementary Dataset 3.

### Hero-cluster NLRs (HCNs) and NRC6 helper genes are nearly exclusively expressed in tomato roots

Given that the potato cyst nematode *Globodera* species and the root-knot nematode *M. enterolobii* that are targeted by the NRC6/HCN resistance gene cluster are root pathogens, we investigated the organ-specific transcriptome profile of these NLR genes. To this end, we performed RNA-sequencing (RNA-seq) on leaf and root tissuess of two-week-old plants of the Hero introgression tomato line LA1792 (Ellis and Maxon Smith, 1971; Ernst et al., 2002). The RNA-seq comparison between leaf and root samples provided consistent data across the three biological replicates. Notably, among the 175 NLR genes with detectable expression, 66% (56% having adjP<0.05) exhibited a root-skewed expression pattern (mean log2FC = 2.05, skewness = 0.49). This contrasted with all expressed tomato genes, where 54% (36% with adjP<0.05) of 28.185 expressed genes showed a root-skewed expression pattern (mean log2FC = 0.23, skewness = 0.03; Supplementary Data 5). This expression pattern of NLR genes can be primarily ascribed to CC-NLRs of the NRC superclade, which display high log2-fold changes for expression in roots (Figure 7A).

**Figure 7.**
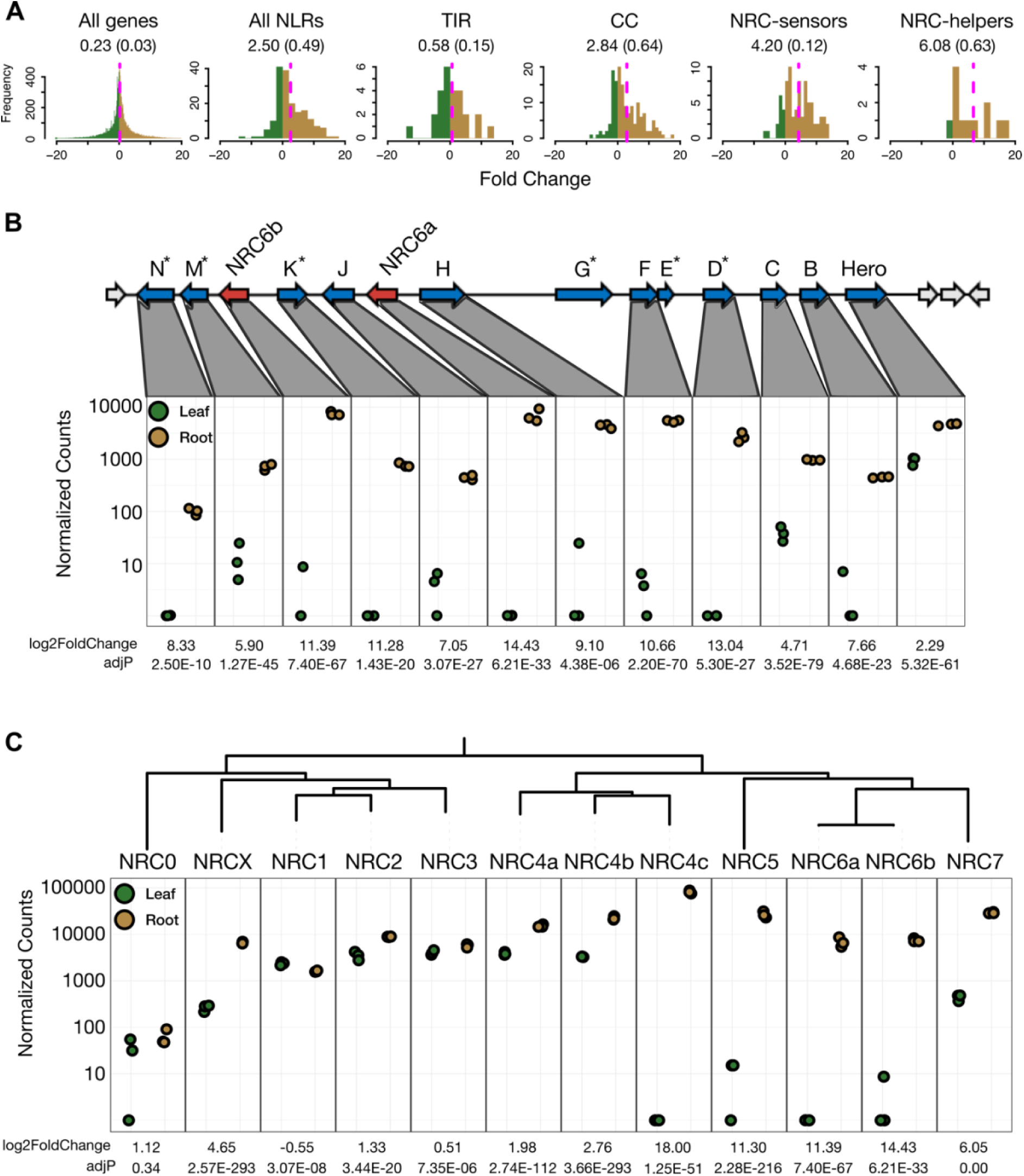
Hero-cluster NLRs (HCNs) and NRC6 helper genes are nearly exclusively expressed in tomato roots. **(A)** Histograms depicting the log2-fold change frequency, comparing the expression levels of all tomato genes, all NLRs, and phylogenetic NLR subclades between roots and leaves of two-week-old unchallenged tomato plants of the Hero introgression line LA1792 (Ellis and Maxon Smith, 1971; Ernst et al., 2002). The dotted magenta line and numbers below labels indicate the mean log2-fold change, the skewness is indicated in brackets. Normalized counts for the expression of cluster-encoded HCN and NRC6 genes **(B)**, and the expression of all tomato NRC clade genes **(C)** in the roots and leaves of two-week-old unchallenged LA1792 tomato plants. The log2-fold change and adjusted p-value are indicated for each gene. Pseudogenes are indicated by an asterisk. HCN-G and HCN-E pseudogenes exhibit negligible expression. Numbers for each gene are presented in Supplementary Table 1. Complete data for log2-fold change values and normalized counts of all tomato genes and NLR subsets are available as Supplementary Data 5; 6.

Next, we examined the 12 genes in the NRC6/HCN gene cluster (Figure 7B). All 10 expressed HCN genes displayed large log2-fold changes for root expression compared to leaves, ranging from 2.28 for Hero to 13.04 for HCN-D (Figure 7B). NRC6a and NRC6b expression was also massively skewed towards roots with log2-fold changes of 14.43 and 11.39, respectively (Figure 7B). Indeed, nine of the 12 genes displayed hardly any RNA-seq signals in leaves, except for HCN-M, HCN-C, and Hero, which showed expression in leaves (Figure 7B, Supplementary Table 1).

Given that *NRC6a* and *NRC6b* are exclusively expressed in roots but not in leaves, we queried our data for the other 10 NRC clade NLRs of tomato (Figure 7C). Except for *NRC0*, *NRC1*, and *NRC3*, the other nine NRCs displayed root-skewed expression patterns with log2-fold changes ranging from 1.33 for *NRC2* to 18 for *NRC4c* (Figure 7C; Supplementary Table 1). Remarkably, besides *NRC6a* and *NRC6b*, *NRC4c* and *NRC5* were also exclusively expressed in roots with hardly any RNA-seq reads observed in leaves, while *NRCX* and *NRC7* are only weakly expressed in leaves compared to roots (Figure 7; Supplementary Table 1). In addition, the strong bias towards root expression can be mapped onto the NRC phylogeny (Figure 7C). The five NRCs in the large clade that comprises *NRC1*, *NRC2*, *NRC3*, and the *NRC4* clade on one hand (three genetically linked paralogs *NRC4a*, *NRC4b* and with the exception of *NRC4c*) show expression in leaves and roots, while the genes in the phylogenetic sister clade comprising *NRC5*, *NRC6a*, *NRC6b* and *NRC7* are exclusively root expressed, or strongly skewed towards root expression (Figure 7C; Supplementary Table 1). In summary, these findings lead us to conclude that the NRC6/HCN gene cluster is exclusively functional in roots, and that root-specific expression is a feature of a number of phylogenetically related NRC helpers in the NRC receptor network.

## Discussion

The NRC network is an intricate immune receptor network that has massively expanded in the lamiid lineage of asterid plants for over 100 million years (Wu et al., 2017; Adachi et al., 2019b; Lee et al., 2021; Goh et al., 2023). However, only a subset of the NRC proteins, which form the network’s central nodes, have been functionally characterized to date. In particular, although genes in the NRC network can be genetically dispersed across the genomes of solanaceous species, such as tomato, a comprehensive analysis of genetic linkage remains outstanding. In addition, there is limited understanding of how clustered and networked NLRs are transcriptionally regulated across different plant organs. In this study, we combined genetic linkage and phylogenetic approaches to identify mixed-clade gene clusters of NLRs in tomato, revealing potential NLR sensor-helper relationships. Our analysis unveiled a mixed-clade gene cluster that is conserved across *Solanum* species and features NRC6 as a helper component and about 10 candidate sensor NLRs (HCNs). Remarkably, all gene cluster-encoded HCNs we tested in this study, including the recently identified root-knot nematode resistance gene *MeR1* and the cyst nematode resistance gene *Hero*, were dependent on *NRC6* for inducing the hypersensitive cell death. Strikingly, none of the other NRC-dependent disease resistance proteins we tested signalled through NRC6, placing this helper NLR in a distinct subnetwork from NRC2, NRC3 and NRC4 (Figure 8). Furthermore, most of the genetically linked *HCNs* and the two *NRC6* paralogs exhibited root-specific expression patterns, suggesting that they form an organ-specific NLR network implicated in resistance against root pathogens (Figure 8). These findings offer novel insights into sub-functionalization within the NRC network, potentially as an adaptive response to organ-specific parasites (Figure 8).

**Figure 8.**
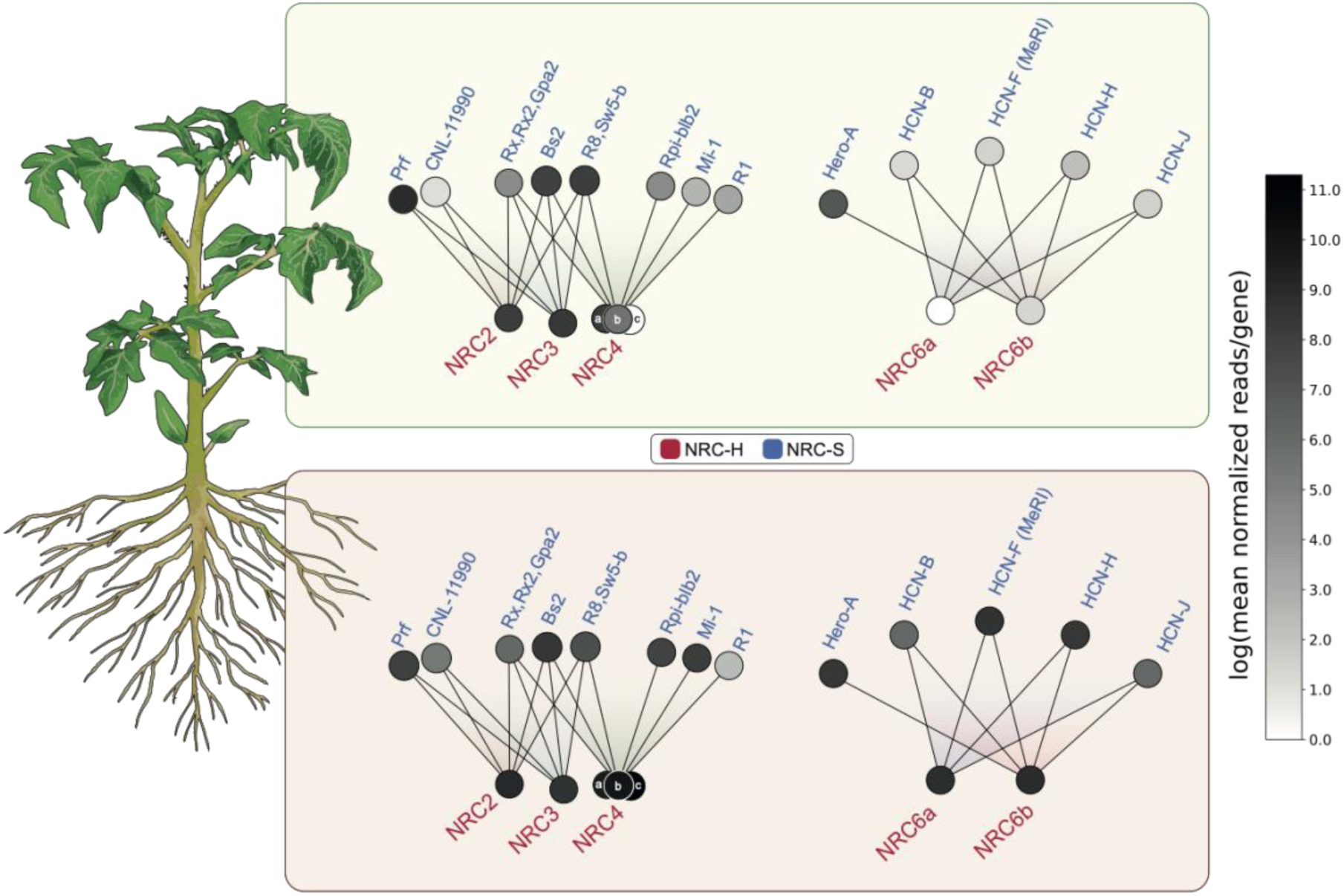
Schematic overview of organ-specific expression patterns in the NRC immune receptor network. Functional connections within the NRC network, involving known sensors and helpers, are represented based on the work of Wu et al., 2017, along with the results presented in this study. Nodes in the network are shaded to reflect their expression levels in either leaves **(top)** or roots **(bottom)**, as indicated by the color scale. Expression levels are based on the mean normalized reads for each gene, derived from RNA-seq data (Supplementary Data 6) obtained from two-week-old unchallenged tomato plants of the Hero introgression line LA1792 (Ellis and Maxon Smith, 1971; Ernst et al., 2002). For NRC sensor genes from other plant species, expression data for the closest tomato homolog is displayed (Supplementary Figure 12).

Our genetic clustering analysis revealed that approximately half of the tomato NLRome is encoded within gene clusters, consistent with previous observations in Arabidopsis (Meyers et al., 2003; Van de Weyer et al., 2019). While most of these gene clusters predominantly contain NLRs from the same phylogenetic clade (Meyers et al., 2003), our analysis identified mixed-clade gene clusters as well (Figure 1, Supplementary Figure 1). In addition to the largest gene cluster containing *NRC6* and *HCNs*, *NRC7* is also linked to an NLR gene from an NRC sensor phylogenetic clade (Figure 1; Figure 2; Supplementary Data 2; Supplementary Dataset 1). Furthermore, the ancestral helper NLR NRC0, genetically clusters with NRC sensors that rely on NRC0 for the induction of cell death (Sakai et al., 2023; Goh et al., 2023; Figure 1; Supplementary Data 2; Supplementary Dataset 1). These findings underscore that mixed-clade gene cluster configurations can be both ancient and relatively recent features of NLR evolution in asterid plants (Sakai et al., 2023; Goh et al., 2023). We propose that unlike the ancient NRC0 gene cluster, the NRC6 and HCN gene cluster has evolved about 17-21 MYA prior to the diversification of the *Solanum* genus, but after the lamiid expansion of the NRC network (Figure 9).

**Figure 9.**
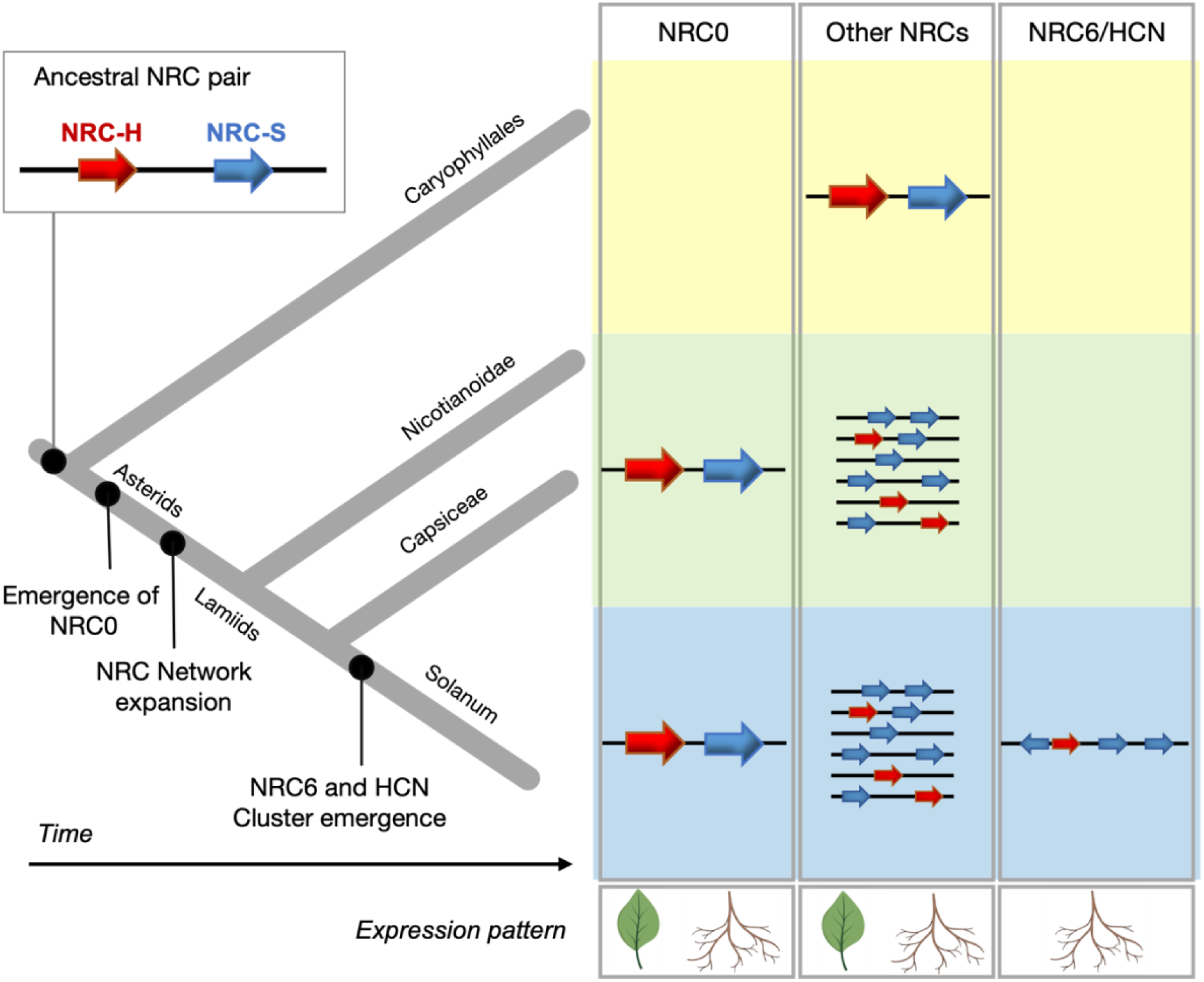
Key steps in the evolution of the NRC network. The ancestral NRC pair is thought to have emerged more than 125 million years ago (MYA) (Sakai *et al*., 2023). We propose that, unlike the ancient NRC0 gene cluster, the NRC6 and HCN gene cluster evolved approximately 17-21 MYA, prior to the diversification of the *Solanum* genus but after the lamiid expansion of the NRC network (Wu et al., 2017; Sakai et al., 2023; Goh et al., 2023). NRC6 and the HCNs formed a gene cluster through genetic duplication and diversification events during the evolution of *Solanum* species. Leaf and root symbols are used to indicate the expression pattern of the respective NLRs.

How did the NRC6/HCN gene cluster emerge? It is plausible that NRC0, previously identified as similar to the ancestral sensor-helper NLR pair of the NRC network found in Asterids and Caryophyllales (Wu et al., 2017; Sakai et al., 2023; Goh et al., 2023), served as the progenitor of the NRC6/HCN gene cluster. This cluster could have emerged following the association of an NRC4/5-derived helper and an SD-type sensor at the same chromosomal location, possibly due to chromosomal or ectopic duplication events. Subsequently, sensor-helper specialization and tandem duplications of sensor NLRs within the gene cluster could have occurred. This scenario is supported by the close phylogenetic relation between NRC6 and NRC4/5, as well as the observation that HCNs have more recently duplicated compared with NRC6 (Seong et al., 2020). While tandem duplication events typically drive cluster expansion, other mechanisms such as unequal crossing-over, gene conversion events, or intra-cluster/gene rearrangements can also contribute to the diversification of NLR gene clusters (Noël et al., 1999; Meyers et al., 2003; Kuang et al., 2004). These mechanisms may provide insights into the wide-ranging diversity of the NRC6 and HCN gene clusters found across various *Solanum* species (Supplementary Figure 3; 4). It is worth noting that reconstructing the sequence of events leading to cluster formation faces significant challenges for NLR genes (Barragan and Weigel, 2021). However, Seong et al. previously identified transposable elements within the NRC6/HCN cluster, suggesting that retroduplications and rearrangements within the cluster could also have shaped its composition (Seong et al., 2020).

Many NRC-dependent sensors form large gene clusters that do not contain an NRC helper (Figure 1; Supplementary Figure 1). An example is the root-knot nematode resistance cluster gene *Mi-1.2* (chromosome 4), which signals through the genetically unlinked helper NRC4 (Wu et al., 2017). *NRC4* forms a cluster on chromosome 4 containing three *NRC4* paralogs (*NRC4a/b/c*; Wu et al., 2020), which are also clustered with *NRC5*, potentially indicating that these NRCs have expanded through tandem duplication events (Figure 1; Supplementary Figure 1B; Wu et al., 2020). Interestingly, the *NRC4* paralogs and *NRC5* are also markedly induced in roots, relative to leaves which coincides with the implicated in resistance against parasitic nematodes similar to the *NRC6* cluster (Figure 7, Supplementary Table 1; Wu et al., 2017). One significant advantage of gene cluster formation through tandem duplications or unequal crossover events is the creation of functionally diverse NLRs (Barragan and Weigel, 2021). We hypothesize that these genomic regions serve as hotspots for generating new resistance genes against root pathogens such as nematodes. This raises the possibility that several uncharacterized nematode resistance genes are encoded in the NRC6/HCN gene cluster as demonstrated by the recent identification of *MeR1* (Frijters et al., 2021). It remains to be tested whether genetically clustered NLRs like *HCNs* and *Mi-1.2* can detect nematode effectors directly or by guarding a host protein targeted by plant-parasitic nematode effectors. The relatively low sequence conservation within the HCN clusters (Seong et al., 2020) may indicate a direct mode of recognition and coevolution of NLRs with pathogen effectors (Contreras et al., 2023b). Recently, a locus of hyper-variable (HYP) effector genes within *G. rostochiensis* and *G. pallida* nematode genomes has been described, potentially harbouring hundreds of allelic effector variants (Sonawala et al., 2023). The existence of such a gene cluster may explain why *Solanum* plant genomes evolved to encode extended, variable clusters of root-expressed NLR genes as a counter-measure.

A long-held view in the field of plant immunity has been that immune receptors are uniformly expressed throughout the entire plant with limited tissue- or cell-specific expression patterns. This perspective may have arisen from the scarcity of expression data in the early days of NLR biology, when many NLR genes were difficult to detect and were poorly annotated. Interestingly, all HCNs and several helper NLRs of the NRC clade exhibited a marked root-skewed baseline expression pattern in tomato (Figure 7; 8, Supplementary Table 1). Indeed, apart from *NRC6*, the phylogenetically related *NRC5* and *NRC7* are strongly expressed in roots (Figure 7). Moreover, *NRC4* paralogs show robust expression in roots (Figure 7; 8, Supplementary Table 1). Several NLRs in the NRC sensor clade also display a root-specific expression pattern, revealing them as potential resistance genes that might signal through root-specific helpers (Figure 7A, Supplementary Figure 11). This observed expression pattern in tomato aligns with the root-skewed NLR expression previously noted in multiple plant species (Gao et al., 2013; Millett et al., 2015; Deng et al., 2017; Munch et al., 2018; Barbey et al., 2019; Adachi et al., 2023). It suggests that the plant immune system may have evolved specialized defences tailored to organ-specific pathogens (Bhardwaj et al., 2011; Wang et al., 2011; Munch et al., 2018; Adachi et al., 2019b; Tang et al., 2023).

Could the organ-specific expression of NLR genes be driven by an evolutionary trend to evade autoimmunity? Elevated expression of NLR genes can potentially harm plant fitness by increasing the risk of unintended activation (Bomblies et al., 2007; Yi and Richards, 2009; Palma et al., 2010). Organ-specific transcriptional regulation is a potential mechanism to mitigate the risk of inadvertent NLR activation, thereby reducing the fitness penalties that may occur when new sensor NLR alleles emerge to detect pathogen effectors. This hypothesis is supported by our observations that some HCNs, such as HCN-B and MeR1, exhibit autoimmunity when co-expressed as wildtype variants with NRC6 in the leaves of *N. benthamiana* (Figure 3, Figure 6). Uncoupling expression between organs or tissues might, for example, offer a means to avoid autoactive cell death in leaves while maintaining baseline expression levels of sensors in roots or other organs targeted by specific pathogens.

How is organ specificity achieved? One way to establish organ-specific expression patterns is through the control of gene expression, using specific transcriptional elements (Dutt et al., 2014). Investigating common promoter motifs, transcription factor binding sites, or the epigenetic regulation of these gene clusters may provide additional insights into the generation of these organ-specific gene expression patterns (Deng et al., 2017; Bartels et al., 2018; Zhou et al., 2022). It is worth noting that NLRs, including those in the tomato NLRome, are also regulated at the transcript level by small RNAs (Zhai et al., 2011). However, our early investigations have not revealed any particularly obvious patterns in the NRC6/HCN cluster genes that would provide cues about root-specific expression. Further research is needed to determine whether the generation of organ- or tissue-specific small RNAs contributes to the observed differences in transcript levels.

The organ-specific facet of NLR biology we documented here has implications for the identification and deployment of disease resistance in agriculture. Considering that plant parasites often infect specific organs and tissues, the expression patterns of NLR genes can accelerate the cloning of novel disease resistance genes by reducing the number of potential candidate genes. Additionally, the development of bioengineered NLRs for resistance against specific pathogens may benefit from organ-specific expression patterns to ensure effective immunity while mitigating the risk of autoimmune responses (Adachi et al., 2019b; Contreras et al., 2023b; Kourelis et al., 2023).

## Materials and Methods

### Gene cluster analysis

Gene cluster analysis was performed as described previously (Sakai et al., 2023). Briefly, the NLR gene annotations were extracted from the ITAG3.2 and RefSeq SL3.0 gff3 files using NLRtracker (Kourelis et al., 2021) and the distance between each NLR gene was determined by a custom script (https://github.com/slt666666/NRC0). NLRs were considered as clustered when the genetic distance was less than or equal to 50 kb. The phylogenetic relationship was based on the NB-ARC domain of extracted sequences and the genetic distance matrix shown in Figure 1 and was generated by using a previously developed “gene-cluster-matrix” library (https://github.com/slt666666/gene-cluster-matrix). All sequences, alignments, phylogenetic tree files, and the interactive matrix file are available as Supplementary Dataset 1.

### Database search for NRC6 and HCN homologs

The full-length NLR sequences retrieved from the NLRtracker (Kourelis et al., 2021) output of 124 Solanaceae genomes (https://doi.org/10.5281/zenodo.10354350) were filtered for sequences with “CNL” and “BCNL” domain architectures. The NB-ARC domain was extracted from the identified 21.833 sequences. Domains shorter than 300 or longer than 400 amino acids were removed as a length distribution parameter applied across all NB-ARC domains. The NB-ARC domains from the RefPlantNLR (Kourelis et al., 2021) dataset, the NRCX dataset (Adachi et al., 2023), HCNs (Hero Cluster NLRs), and tomato NRCs were incorporated, resulting in a refined dataset containing 20.292 sequences. The NB-ARC domains were aligned using MAFFT v7.505 (Katoh et al., 2002; Katoh and Standley, 2013) with the [--anysymbol] option, and a phylogenetic tree was constructed using FastTree v2.1.10 (Price et al., 2010) with the [-lg] option. A custom R script was employed for sub setting of well-supported major branches to generate specific sub-trees for NRC helper, NRC-SD sensor, and HCN clades. The resulting phylogenetic trees were visualized using iTOL (Letunic and Bork, 2021) and manually annotated. All scripts utilized in this analysis are deposited under https://github.com/amiralito/Hero. All sequences, alignments and phylogenetic tree files are available as Supplementary Dataset 2.

### Determination of NRC6 and HCN gene clusters from genome-scale assemblies of Solanum plants

BLAST (Altschul et al., 1990) searches for Hero and NRC6 homologs were performed in Geneious Prime (https://www.geneious.com), against local databases of genome sequences and annotations of wild tomato (*S. pennelli*, SPENNV200, Bolger et al., 2014), potato (*S. tuberosum*, DM 1-3 516 R44, Potato Genome Sequencing Consortium et al., 2011), and American black nightshade (*S. Americanum*, SP2271, Lin et al., 2023). Genes in the genomic loci of the received top hits were extracted, translated, and annotated to verify the presence of an NB-ARC domain (Pfam ID: PF00931). The detected NLR gene clusters were drawn to scale.

### Plant growth condition

Wildtype and *nrc2/3/4* mutant (Wu et al., 2020) *N. benthamiana* plants were grown in a glasshouse for all cell death assays performed. Wildtype *N. benthamiana* plants used for protein extraction, and the Hero tomato line (*S. lycopersicum*, LA1792) used for RNA-extraction and RNA-seq were grown in a growth chamber at 22–25°C, 45–65% humidity, and a 16/8 hr light/dark cycle.

### Plasmid construction

The Golden Gate Modular Cloning (MoClo; Weber et al., 2011) and MoClo plant parts kits (Engler et al., 2014) were used for cloning. Wildtype and MHD-mutant variants of tomato NRC helpers and HCNs were synthesized as *N. benthamiana* codon-optimized L0 modules in pICH41155 through GENEWIZ/AZENTA (https://www.azenta.com/). All plasmids generated in this study were cloned into the binary vector pJK001c (Paulus et al., 2020). Cloning design and sequence analysis were performed in Geneious Prime (https://www.geneious.com). All Plasmids used and constructed in this study are described in Supplementary Data 7.

### Transient gene expression and cell death assays

Transient gene expression in *N. benthamiana* was performed by agroinfiltration. *A. tumefaciens* strain GV3101 pMP90 transformed with respective binary expression constructs were inoculated from glycerol stocks and grown O/N at 28°C in LB supplemented with appropriate antibiotics. Cells were harvested by centrifugation at 2000 × g for 10 min at RT and resuspended in infiltration buffer (10 mM MgCl_2_, 10 mM MES-KOH pH 5.6, 200 μM acetosyringone). Cells were left to incubate in the dark for 2 hours at room temperature prior to infiltration into 5 to 6-week-old *N. benthamiana* leaves at ODs indicated in Supplementary Data 7. Cell death phenotypes were scored with a range from 0 (no visible necrosis) to 7 (fully confluent necrosis) according to Adachi et al., 2019a). Quantification and statistical analysis was performed by using the besthr R library (MacLean, 2019) and plotted using a script described in Bentham et al., 2023. Scoring for all experiments can be found in Supplementary Dataset 4.

### Protein extraction and SDS-PAGE assay

*N. benthamiana* leaf discs (8 mm diameter) were taken 2 days post-infiltration (dpi) with Agrobacteria and were homogenized in extraction buffer [10% glycerol, 25 mM Tris-HCl, pH 7.5, 1 mM EDTA, 150 mM NaCl, 1% (w/v) PVPP, 10 mM DTT, 1x protease inhibitor cocktail (SIGMA), 0.2% IGEPAL CA-630 (SIGMA)]. After centrifugation at 12,000 x g for 10 min at 4°C, the obtained supernatant was mixed with 2x SDS loading buffer [final concentration: 50 mM Tris-HCl (pH 6.8), 100 mM DTT, 2% SDS, 0.01% bromophenol blue, 10% glycerol] and denatured at 72°C for 10 min. Total protein extracts were separated by SDS-PAGE gels (Bio-Rad) and transferred onto polyvinylidene difluoride (PVDF) membranes using a Trans-Blot turbo transfer system (BioRad). Membranes were blocked for 60 min in 5% milk powder dissolved in Tris-buffered Saline [50mM Tris-HCL (pH7.5), 150mM NaCl]. Mouse monoclonal anti-GFP antibody conjugated to HRP (B-2, Santa Cruz Biotech; 1:5000 dilution) or mouse monoclonal anti-mCherry TrueMAB antibody conjugated to HRP (OTI10G6, Thermo Fisher Scientific; 1:2500 dilution) were used to probe the membranes. Equal loading was monitored by staining the PVDF membranes with Ponceau S (SIGMA).

### RNA-extraction, RNA-seq analysis and HCN reading frame adjustment

Total RNA from leaf and root tissues of two-week-old *S. lycopersicum*, LA1792 was extracted from independent biological triplicates using the RNeasy Mini Kit (Qiagen). Each sample was sent for library preparation and Illumina NovaSeq 6000 sequencing (40 M paired-end reads per sample, Novogene). High-quality reads were mapped to a pseudogenome generated by replacing the NRC6 and HCN gene cluster of the Heinz reference assembly (SL3.0, ITAG3.2 annotation) with the cluster sequence obtained by Ernst et al., 2002, using HISAT2 (Kim et al., 2019). Read alignment to the NRC6 and HCN gene cluster was used to correct reading frames and sequences of NRC and HCN gene cluster encoded genes as an iterative approach. The CDS sequences of HCN-B (OR865987), HCN-C (OR865986), HCN-F (OR865983), HCN-H (OR865981), HCN-J (OR865982), NRC6a (OR865985), and NRC6b (OR865984) were deposited to the National Center for Biotechnology Information (NCBI) GenBank. Read counting was performed using STRINGTIE2 (Pertea et al., 2015; Kovaka et al., 2019) and differential expression of genes was determined by DeSeq2 (Love et al., 2014), comparing leaf against root tissues. Raw reads used in this study were deposited in the NCBI Sequence Read Archive (SRA) with BioSample accession SAMN38499990 and BioProject accession PRJNA1046475. The pseudogenome, pseudoannotation, original NRC6/HCN cluster sequences generated by Ernst et al., 2002, as well as the corrected cluster sequences and annotations are available under https://zenodo.org/records/10376142. Scripts used for RNA-seq analysis are deposited under https://github.com/amiralito/Hero.

## Author contributions

Conceptualization: S.K., D.L., H.A., C.H.W., J.K.,

Data Curation: D.L., T.S., A.T., J.K., A.P., C.H.W

Formal Analysis: D.L., T.S., A.T., J.K., A.P, C.H.W., A.V., R.F., K.E., M.G.,

Investigation: D.L., T.S., A.T., J.K., A.P, H.P., A.H., C.H.W., A.V., R.F, K.E.

Methodology: D.L., T.S., A.T., J.K., A.P, H.P., A.H., C.H.W., A.V., R.F, K.E.

Resources: D.L., T.S., A.T., J.K., A.P, H.P., A.H., C.H.W., A.V., R.F, K.E.

Software: T.S., A.T., A.P., D.L

Supervision: S.K., H.A., C.H.W.

Funding Acquisition: S.K.

Project Administration: S.K.

Writing Initial Draft: D.L., S.K.

Editing: D.L., S.K., J.K., A.T., C.H.W., H.A.

## Acknowledgments

We are thankful to our Sainsbury Laboratory colleagues Mauricio P. Contreras, Clemence Marchal, Cristina Barragan, Thorsten Langner, and Joe Win for helpful discussions, ideas, and support. This work was funded by the Gatsby Charitable Foundation, Biotechnology and Biological Sciences Research Council (BBSRC, UK, BB/WW002221/1, BB/V002937/1, BBS/E/J/000PR9795 and BBS/E/J/000PR9796) and the European Research Council (BLASTOFF). J.K. was funded by BASF Plant Science. H.A. was funded by the Japan Science and Technology Agency, Precursory Research for Embryonic Science and Technology (JPMJPR21D1). C.H.W. was funded by the 2030 Cross-Generation Young Scholars Program of the National Science and Technology Council, Taiwan (NSTC 112-2628-B-001-007). K. E. was funded by the DFG grant Ga470/1/1-3. D.L. was funded by the DFG Walter Benjamin Programme—project no. 464864389. The funders had no role in the preparation of the manuscript.

## Supplementary Figures

**Supplementary Figure 1.**
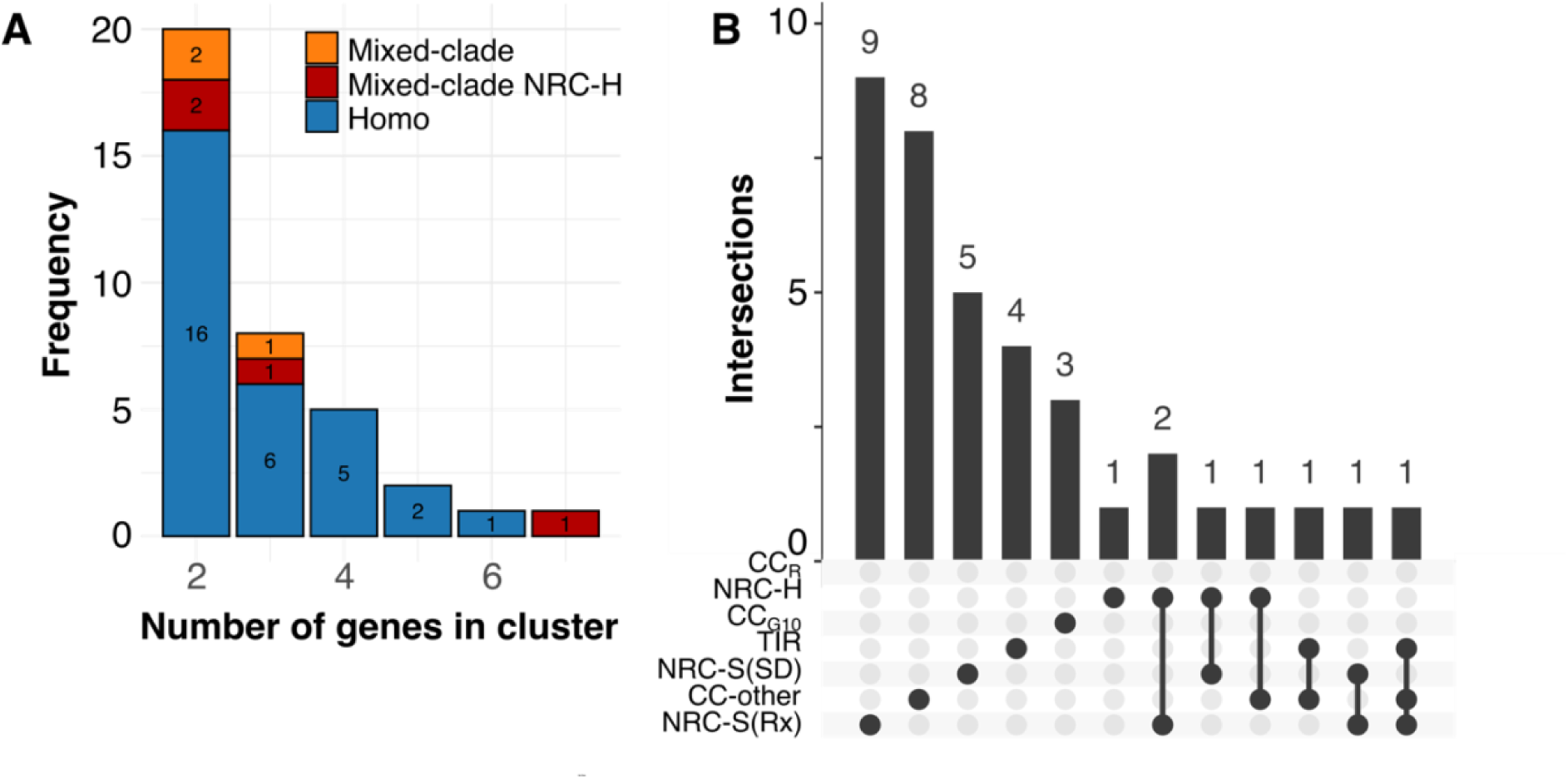
The tomato genome encodes homo and mixed-clade NLR gene clusters with different compositions. (A) Frequency of NLR gene cluster sizes found in the tomato genome. The number of homo, mixed-clade gene clusters, and mixed-clade gene clusters containing NRC helpers are indicated. (B) Upset plot showing the frequency for cluster compositions for all detected tomato NLR gene clusters. The frequency of homo clusters for respective NLR clades are indicated by a single dot, frequency of mixed-clade gene clusters and the respective composition is indicated by dots joined together by a line.

**Supplementary Figure 2.**
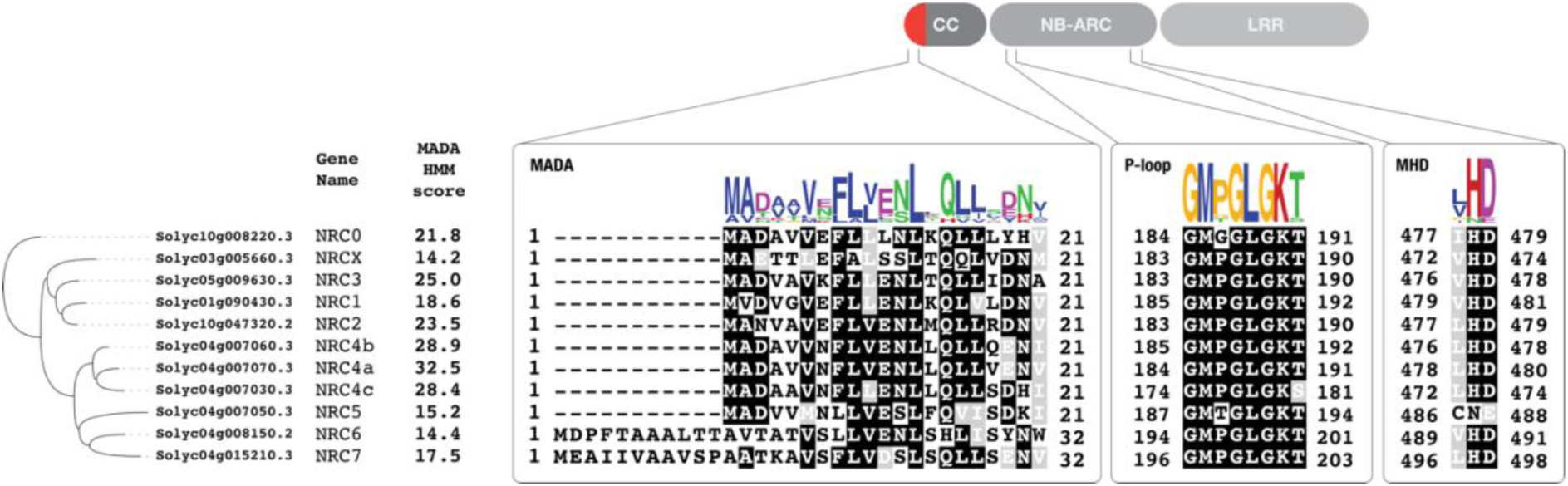
NRC6 is a canonical NRC helper with an extension before the MADA motif. Schematic representation of the NRC helper domain architecture (top), depicting CC, NB-ARC, and LRR domains. The position of the α1 helix within the CC domain, containing the MADA motif, is indicated in red. The MADA motif score, based on a Hidden-Markov model (Adachi et al., 2019), is shown for each tomato NRC helper. A MEME motif was generated for the MADA motif, the P-loop motif, and the MHD motif based on the amino acid sequence alignment of all tomato NRC helpers. The respective amino acid positions of the motifs within each NRC helper are indicated.

**Supplementary Figure 3.**
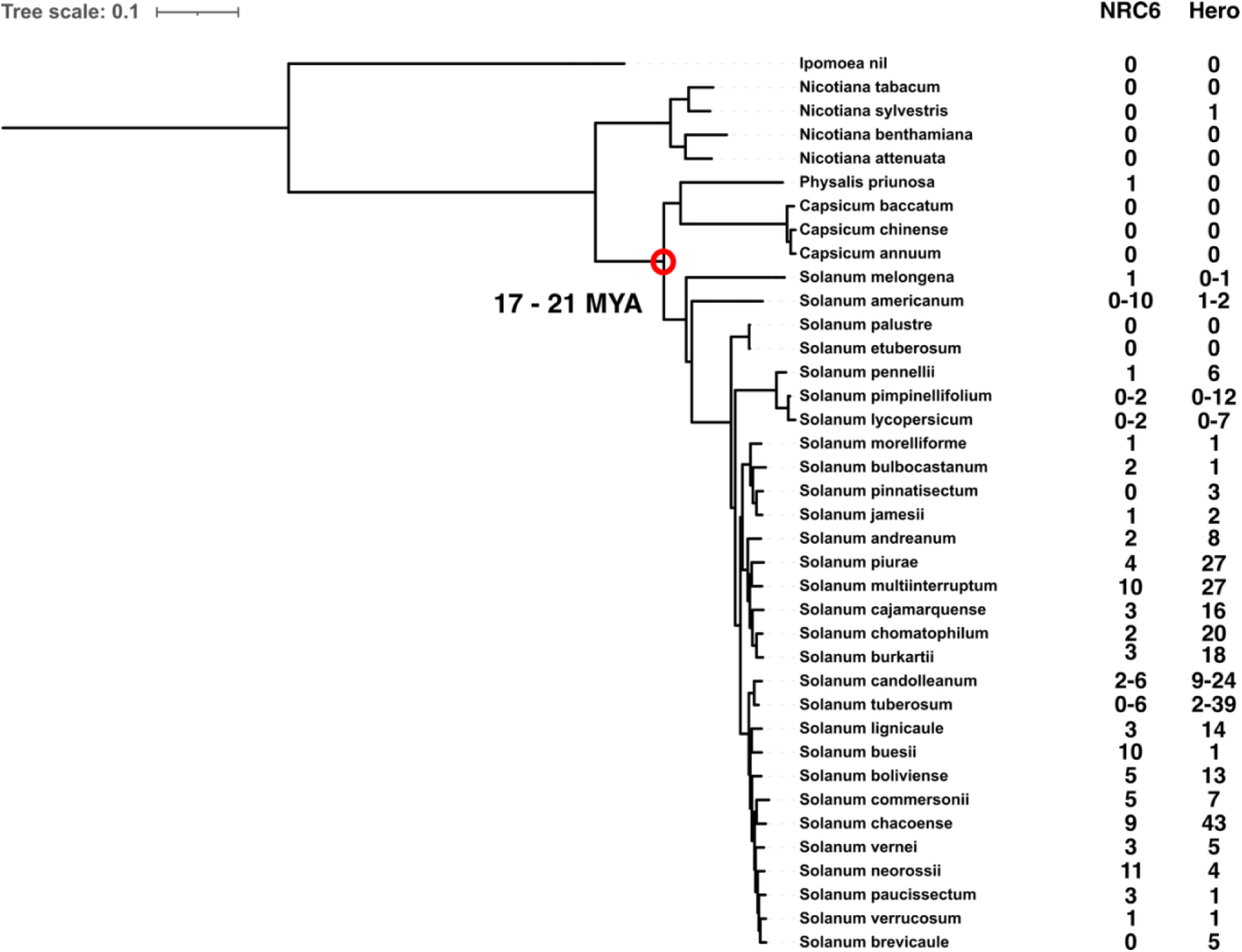
The NRC6/HCN gene cluster emerged in Solanum plants. A phylogenetic tree illustrating the species relationships between Caryophyllales and asterid plant species is presented. NRC6, along with Hero-homologs, is exclusively encoded within Solanum plants (Supplementary Data 4). The range of numbers for NRC6 and Hero-homologs in each species is based on phylogeny, using the data displayed in Figure 2 and Supplementary Dataset 2. The species phylogeny was derived from (Wu et al., 2023), the time of divergence is based on (Särkinen et al., 2013).

**Supplementary Figure 4.**
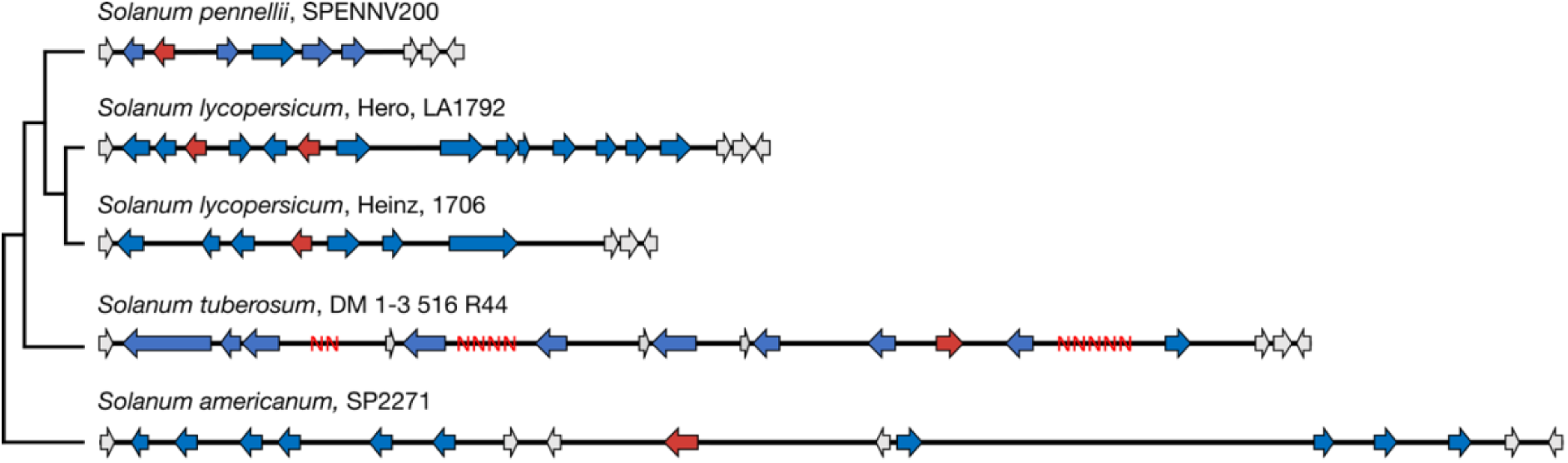
Comparison of the NRC6/HCN gene cluster in Solanum species. Schematic representation of the NRC6 and HCN gene cluster extracted from chromosome-scale genome sequence assemblies of wild tomato (*S. pennelli*), potato (*S. tuberosum*), and American black nightshade (*S. americanum*), together with the gene clusters of *S. lycopersicum* Heinz 1706 and Hero LA1792, which contains an introgressed cluster from *S. pimpinellifolium* LA121 (Ellis and Maxon Smith, 1971; Ernst et al., 2002). The NRC6 helper is colored in red, and HCNs are colored in blue. Gaps in the sequence assembly are depicted as red Ns.

**Supplementary Figure 5.**
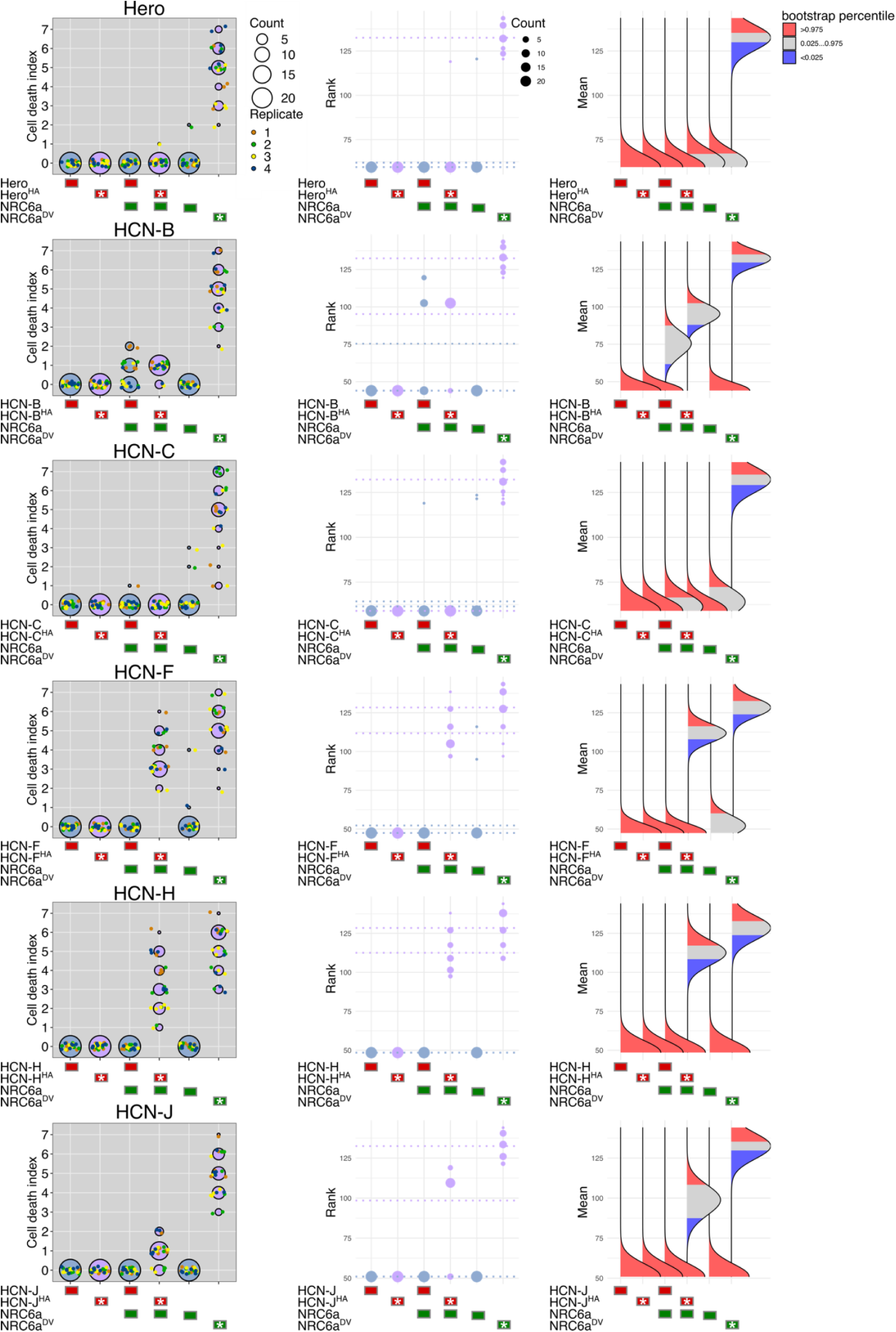
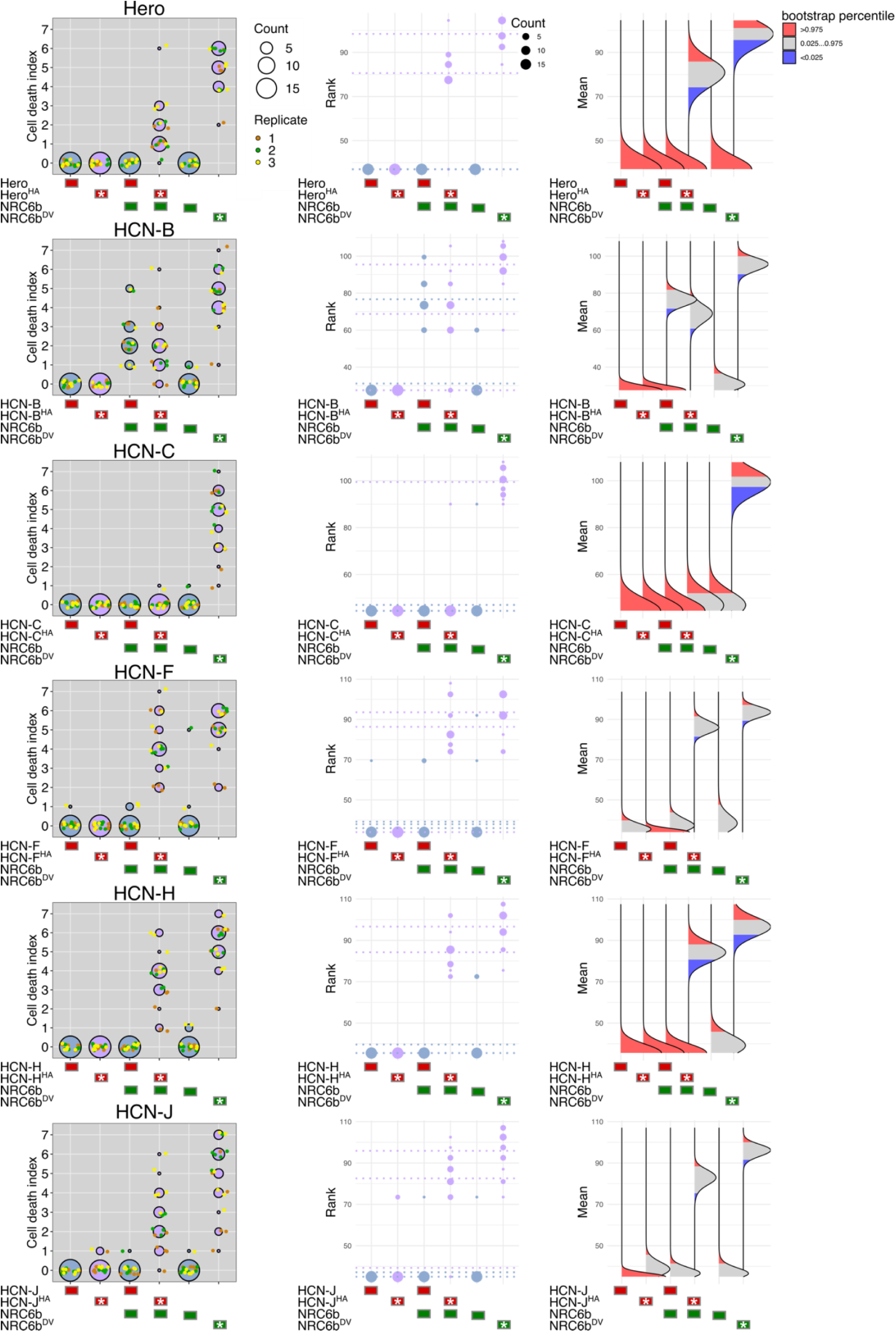
Quantification and statistical analysis of cell death upon co-expression of HCNs with NRC6a or NRC6b. (left) Cell death data represented as dots, colored for each biological replicate. The central circle for each cell death category proportionally represents the total number of data points. An asterisk indicates expression of an autoactive NLR mutant (autoactive HCN = HA; autoactive NRC6b = DV). **(middle)** Statistical analysis using the besthr R library (MacLean, 2019). The ranked data is shown as dots with their corresponding mean as dashed line. The size of each dot proportionally represents the total number of data points. A bootstrap resampling test, using a lower significance cutoff of 0.025 and an upper cutoff of 0.975, was performed. Mean ranks of test samples falling outside of these cutoffs in the control samples bootstrap population (respective HCN alone) were considered significant. **(right)** The distribution of 1,000 bootstrap sample rank means, blue areas under the curve illustrate the 0.025, and red areas the 0.975 percentiles of the distribution.

**Supplementary Figure 6.**
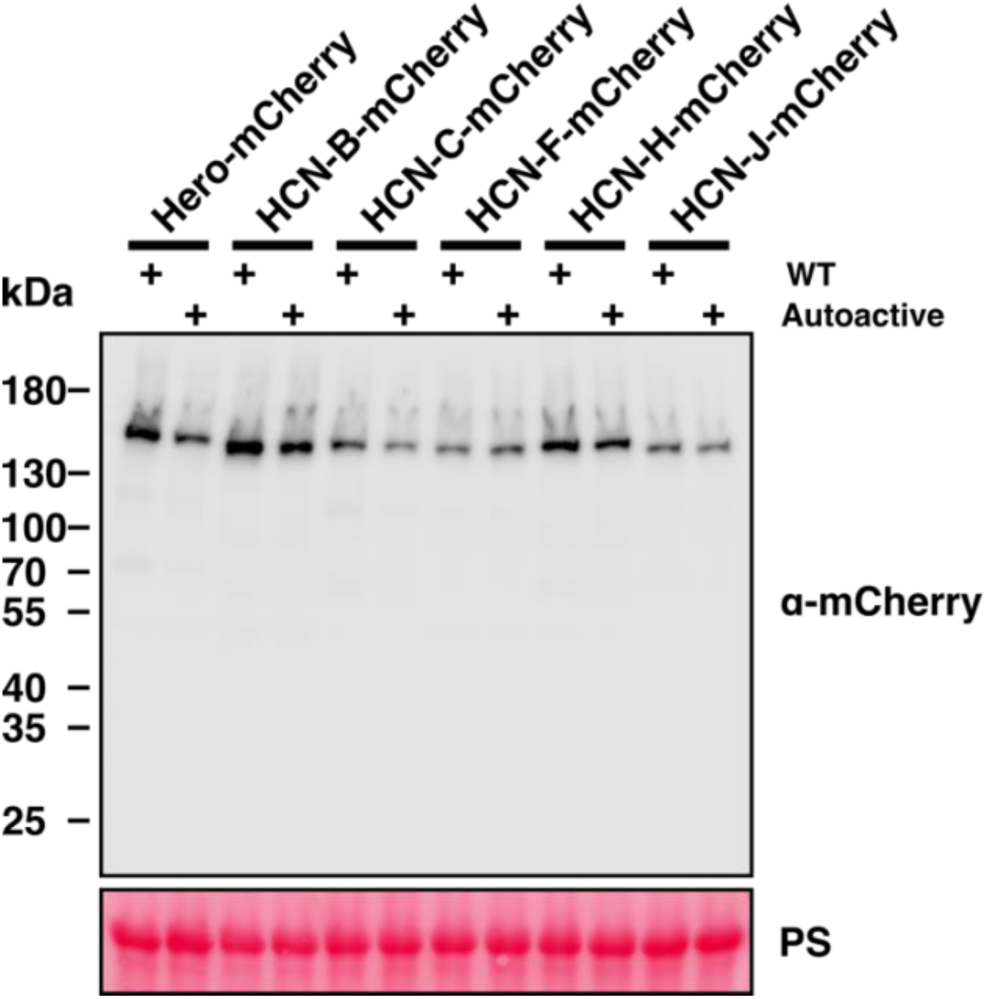
Accumulation of wildtype and autoactive HCNs in planta. Immunoblot analysis depicting the accumulation of HCN-mCherry in wildtype *N. benthamiana* plants at 2 days post-infiltration (dpi) with Agrobacteria transformed with the respective expression constructs. PS, Ponceau S stain.

**Supplementary Figure 7.**
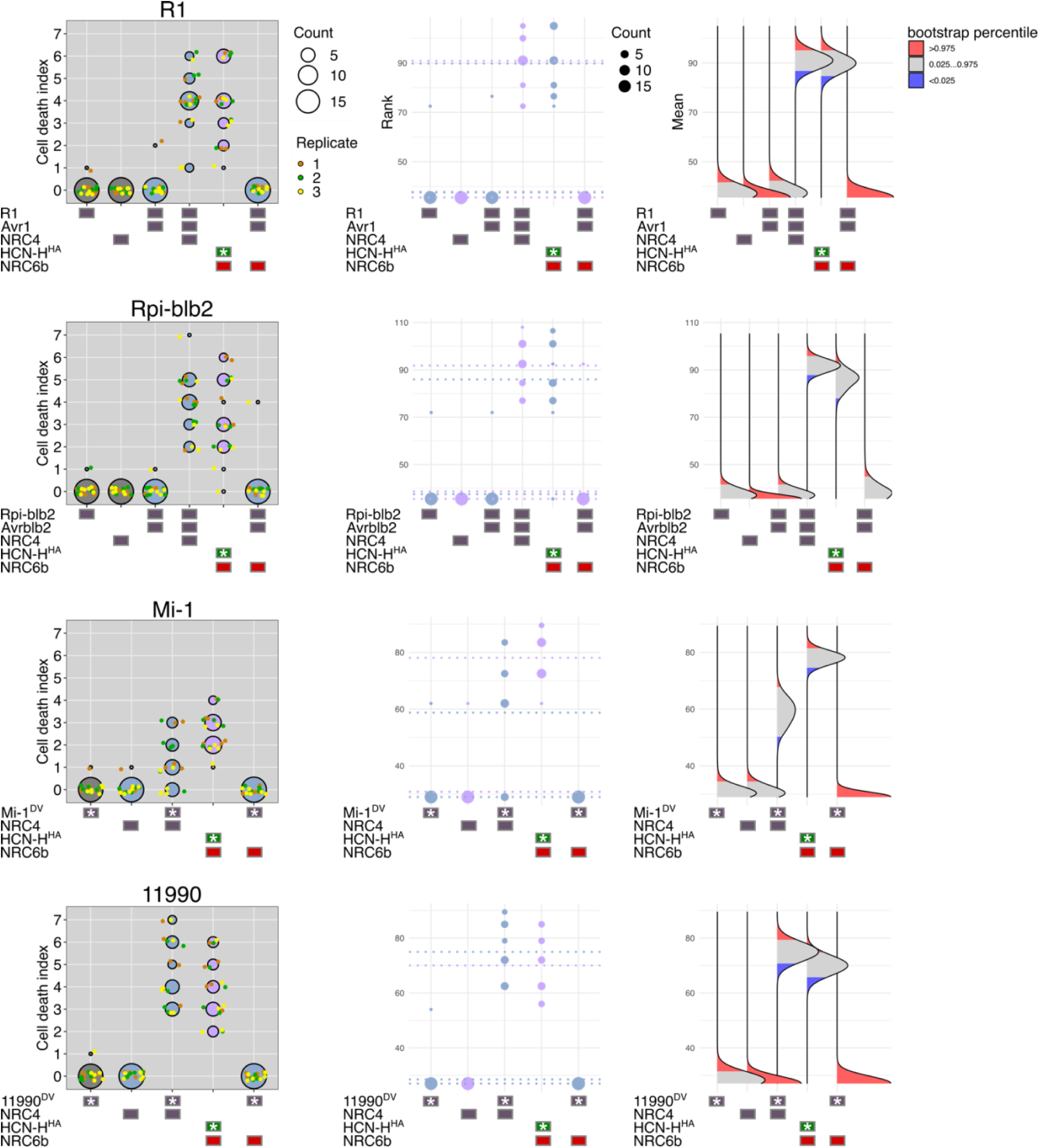
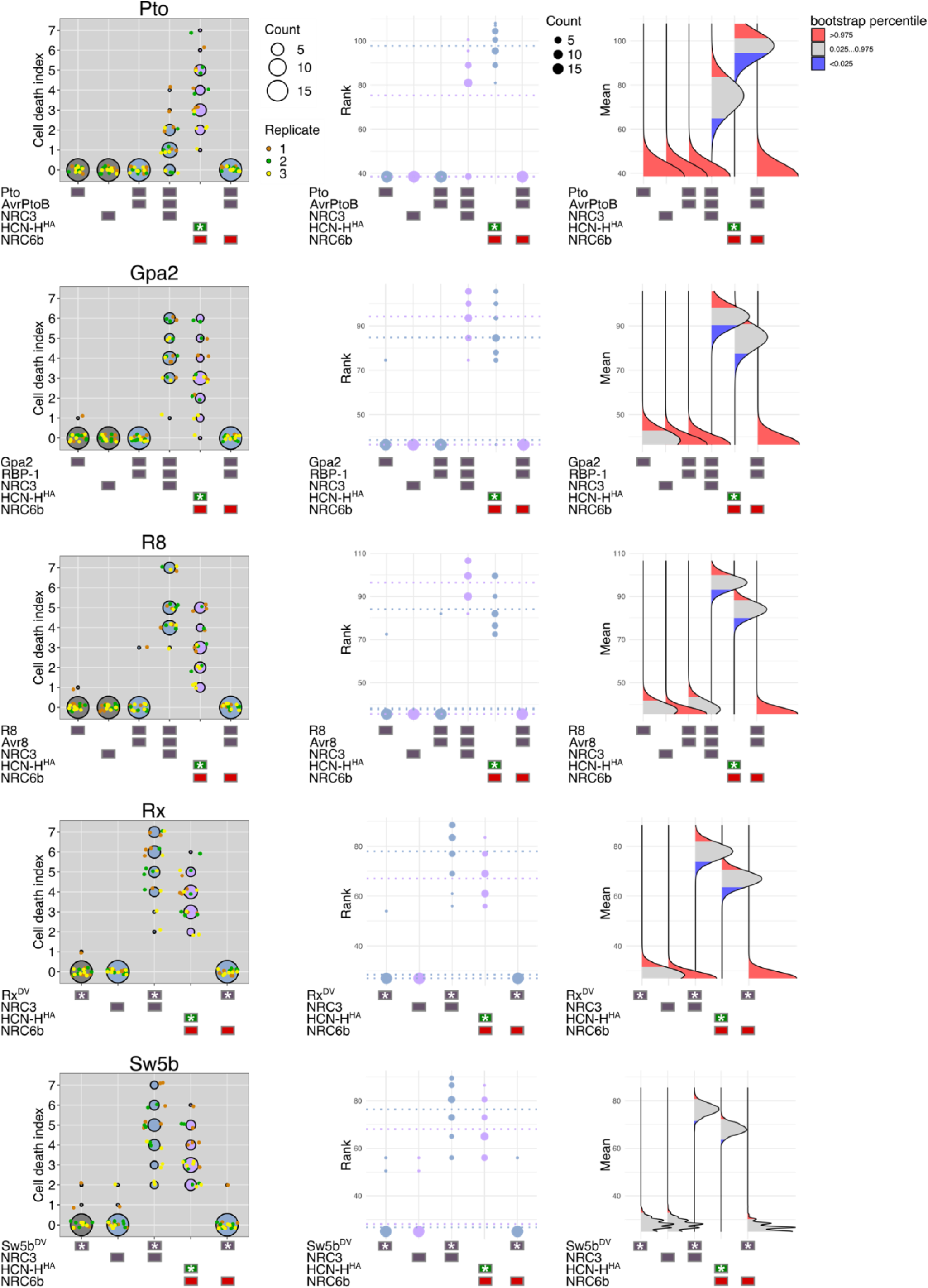
Quantification and statistical analysis of cell death upon co-expression of NRC2/3/4-dependent sensors. (left) Cell death data represented as dots, colored for each biological replicate. The central circle for each cell death category proportionally represents the total number of data points. An asterisk indicates expression of an autoactive NLR mutant. **(middle)** Statistical analysis using the besthr R library (MacLean, 2019). The ranked data is shown as dots with their corresponding mean as dashed line. The size of each dot proportionally represents the total number of data points. A bootstrap resampling test, using a lower significance cutoff of 0.025 and an upper cutoff of 0.975, was performed. Mean ranks of test samples falling outside of these cutoffs in the control samples bootstrap population (sensor alone) were considered significant. **(right)** The distribution of 1,000 bootstrap sample rank means, blue areas under the curve illustrate the 0.025, and red areas the 0.975 percentiles of the distribution.

**Supplementary Figure 8.**
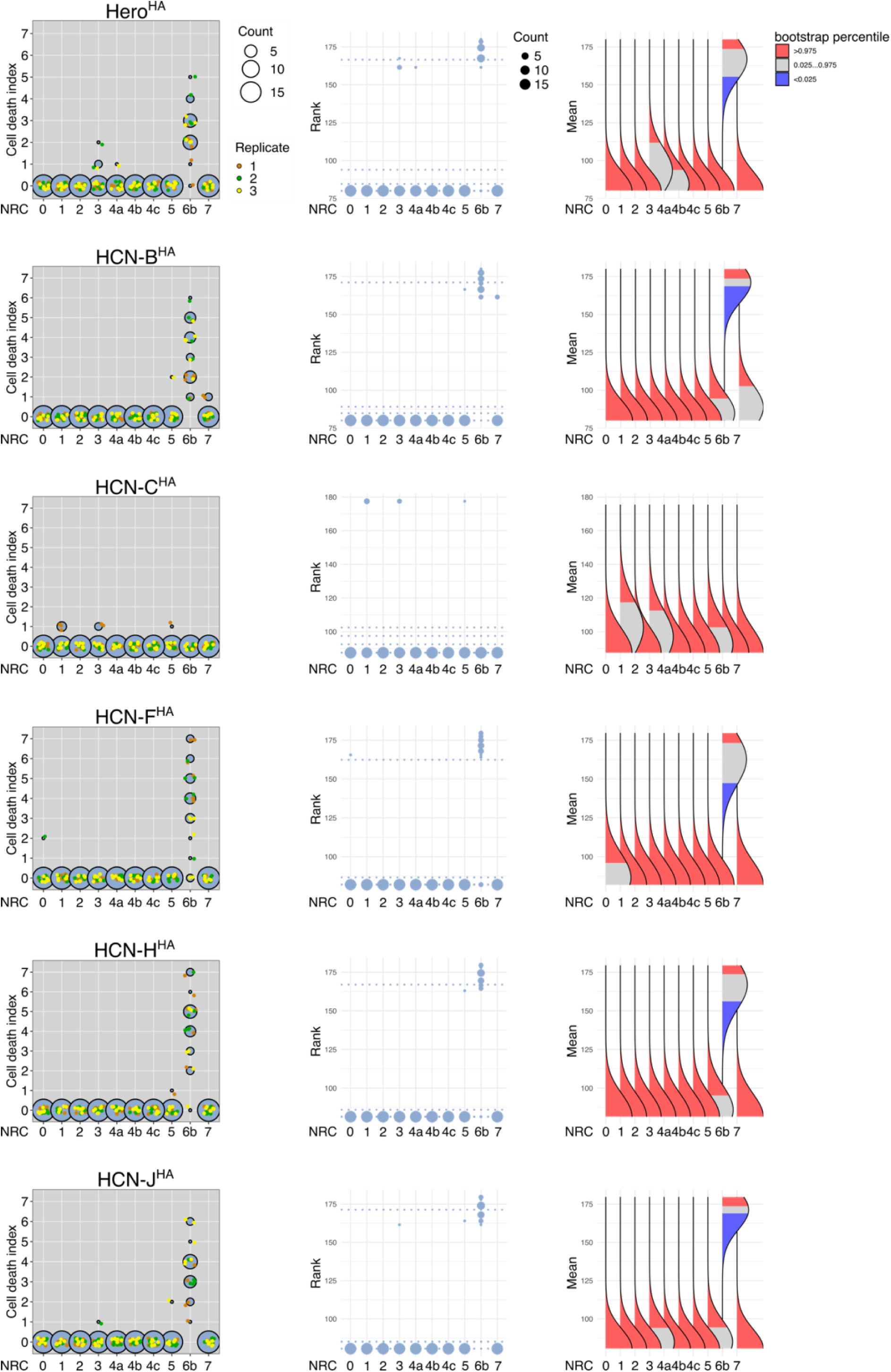
Quantification and statistical analysis of cell death upon co-expression of autoactive HCNs with tomato NRCs. (left) Cell death data represented as dots, colored for each biological replicate. The central circle for each cell death category proportionally represents the total number of data points. An asterisk indicates expression of an autoactive NLR mutant. **(middle)** Statistical analysis using the besthr R library (MacLean, 2019). The ranked data is shown as dots with their corresponding mean as dashed line. The size of each dot proportionally represents the total number of data points. A bootstrap resampling test, using a lower significance cutoff of 0.025 and an upper cutoff of 0.975 was performed. Mean ranks of test samples falling outside of these cutoffs in the control samples bootstrap population (NRC0) were considered significant. **(right)** The distribution of 1,000 bootstrap sample rank means, blue areas under the curve illustrate the 0.025, and red areas the 0.975 percentiles of the distribution.

**Supplementary Figure 9.**
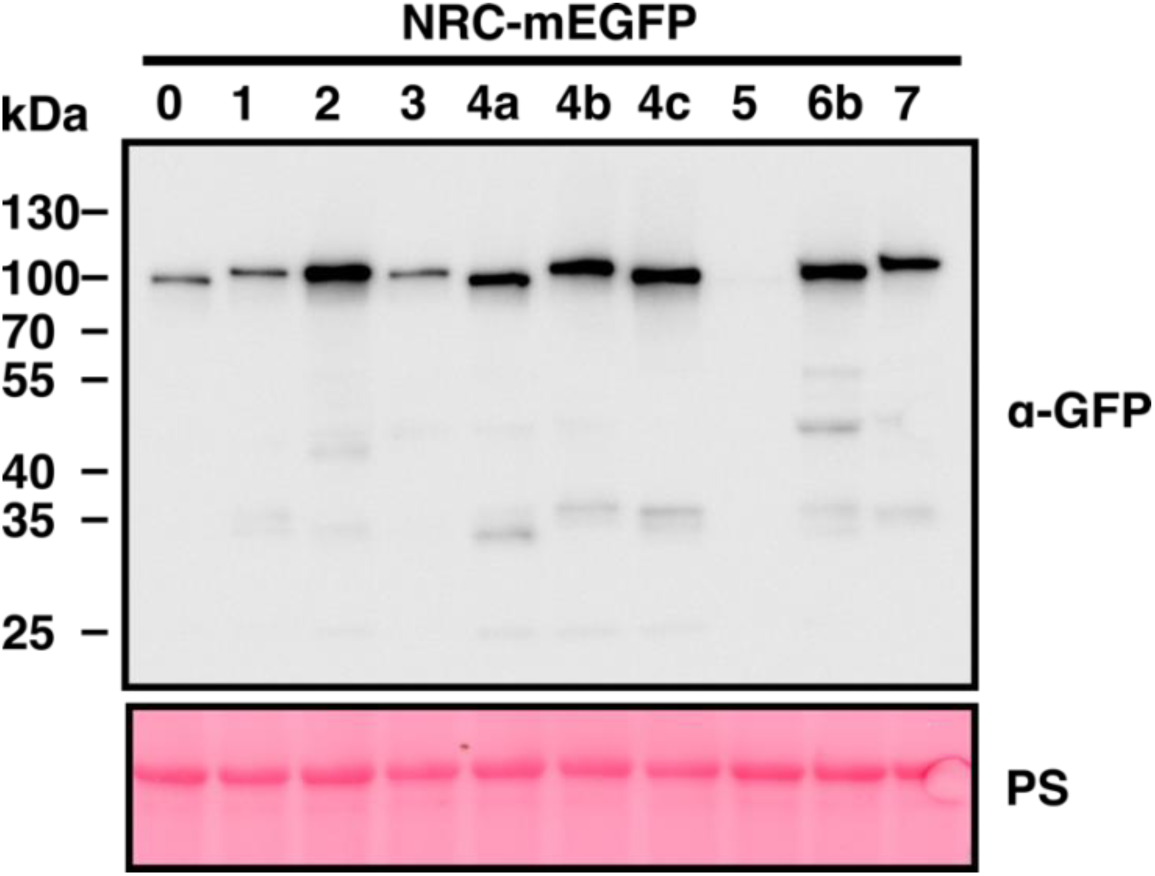
Tomato NRC helpers accumulate in planta. Immunoblot analysis of NRC mEGFP accumulation in *N. benthamiana nrc2/3/4* mutant plants at 2 days post-infiltration (dpi) with Agrobacteria transformed with the respective expression constructs. PS, Ponceau S stain.

**Supplementary Figure 10.**
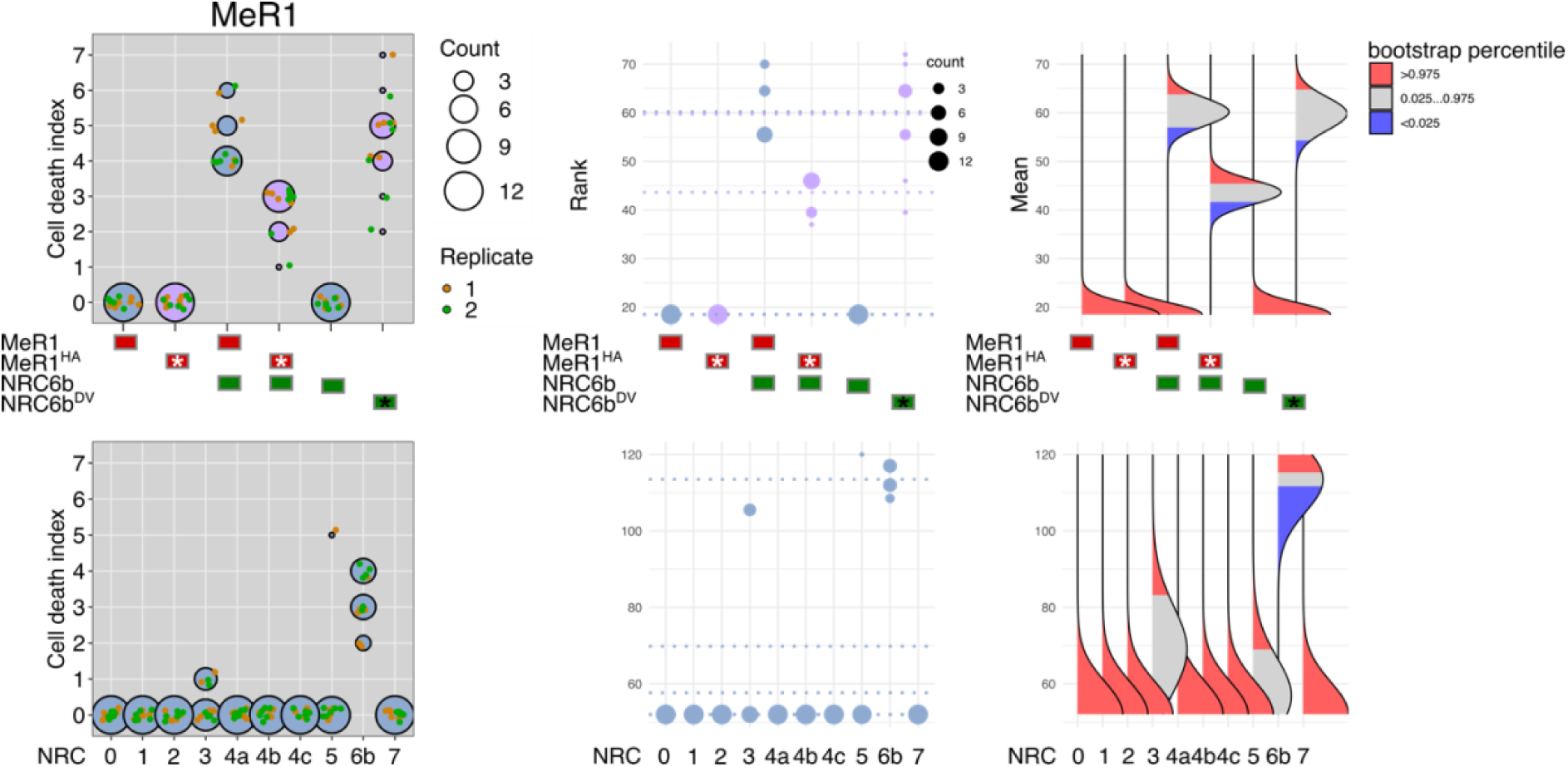
Quantification and statistical analysis of cell death upon co-expression of wildtype or autoactive MeR1 with NRC6 or other tomato NRC helpers. (left) Cell death data represented as dots, colored for each biological replicate. The central circle for each cell death category proportionally represents the total number of data points. An asterisk indicates expression of an autoactive NLR mutant. **(middle)** Statistical analysis using the besthr R library (MacLean, 2019). The ranked data is shown as dots with their corresponding mean as dashed line. The size of each dot proportionally represents the total number of data points. A bootstrap resampling test, using a lower significance cutoff of 0.025 and an upper cutoff of 0.975 was performed. Mean ranks of test samples falling outside of these cutoffs in the control samples bootstrap population (MeR1 alone (top) or NRC0 (bottom)) were considered significant. **(right)** The distribution of 1,000 bootstrap sample rank means, blue areas under the curve illustrate the 0.025, and red areas the 0.975 percentiles of the distribution.

**Supplementary Table 1.** Expression and log2-fold change of HCN and NRC genes in tomato roots and leaves. The expression of genes is given as normalized counts for each independent biological sample of leaves (L1, L2, L3) and roots (R1, R2, R3) of two-week-old unchallenged LA1792 tomato plants, with the corresponding log2-fold change (log2FC) and adjusted P-value (adjP). Expression data and log2-fold changes for all tomato genes is shown in Supplementary Data 5 and 6.

| Gene | L1 | L2 | L3 | R1 | R2 | R3 | log2FC | adjP |
| --- | --- | --- | --- | --- | --- | --- | --- | --- |
| Hero | 1046.31 | 1032.22 | 762.56 | 4729.93 | 4810.30 | 4365.15 | 2.29 | 7.32E-61 |
| HCN-B | 6.53 | 0.00 | 0.00 | 452.44 | 464.87 | 435.89 | 7.66 | 4.68E-23 |
| HCN-C | 26.15 | 36.76 | 50.43 | 947.75 | 997.42 | 969.90 | 4.71 | 3.52E-79 |
| HCN-D | 0.00 | 0.00 | 0.00 | 2452.05 | 3337.51 | 2098.27 | 13.04 | 5.30E-27 |
| HCN-F | 3.26 | 5.88 | 0.00 | 5528.69 | 5588.93 | 5158.00 | 10.66 | 2.20E-70 |
| HCN-H | 0.00 | 0.00 | 24.20 | 4639.44 | 3896.83 | 4546.73 | 9.10 | 4.38E-06 |
| HCN-J | 0.00 | 5.88 | 4.03 | 406.18 | 434.96 | 491.72 | 7.05 | 3.07E-27 |
| HCN-K | 0.00 | 0.00 | 0.00 | 854.54 | 734.55 | 726.86 | 11.28 | 1.43E-20 |
| HCN-M | 9.80 | 4.41 | 24.20 | 621.17 | 746.10 | 788.89 | 5.90 | 6.27E-45 |
| HCN-N | 0.00 | 0.00 | 0.00 | 83.00 | 104.93 | 113.34 | 8.33 | 2.50E-10 |
| NRC0 | 31.06 | 54.40 | 0.00 | 48.30 | 90.77 | 47.36 | 1.12 | 0.34 |
| NRCX | 214.16 | 282.31 | 292.51 | 6381.87 | 6936.32 | 6442.56 | 4.65 | 2.57E-293 |
| NRC1 | 2516.05 | 2371.75 | 2178.76 | 1598.87 | 1568.28 | 1672.53 | -0.55 | 3.07E-08 |
| NRC2 | 4195.06 | 3559.83 | 2785.98 | 8738.68 | 8874.51 | 8966.58 | 1.33 | 3.44E-20 |
| NRC3 | 3655.56 | 3923.03 | 4502.77 | 5900.17 | 6118.35 | 5227.93 | 0.51 | 7.35E-06 |
| NRC4a | 4172.17 | 3631.89 | 3788.62 | 16352.71 | 14693.26 | 14601.09 | 1.98 | 2.74E-112 |
| NRC4b | 3264.82 | 3333.40 | 3324.63 | 24577.03 | 20867.75 | 21654.92 | 2.76 | 3.66E-236 |
| NRC4c | 0.00 | 0.00 | 0.00 | 86660.85 | 75688.36 | 81911.12 | 18.00 | 1.25E-51 |
| NRC5 | 14.71 | 14.70 | 0.00 | 31201.80 | 23106.58 | 25870.65 | 11.30 | 2.28E-216 |
| NRC6a | 0.00 | 0.00 | 0.00 | 8852.98 | 5469.31 | 6324.71 | 11.39 | 7.40E-67 |
| NRC6b | 8.17 | 0.00 | 0.00 | 8243.37 | 7128.36 | 7111.35 | 14.43 | 6.21E-33 |
| NRC7 | 480.65 | 367.60 | 484.17 | 30304.39 | 28525.00 | 28782.07 | 6.05 | 0.00 |

**Supplementary Figure 11.**
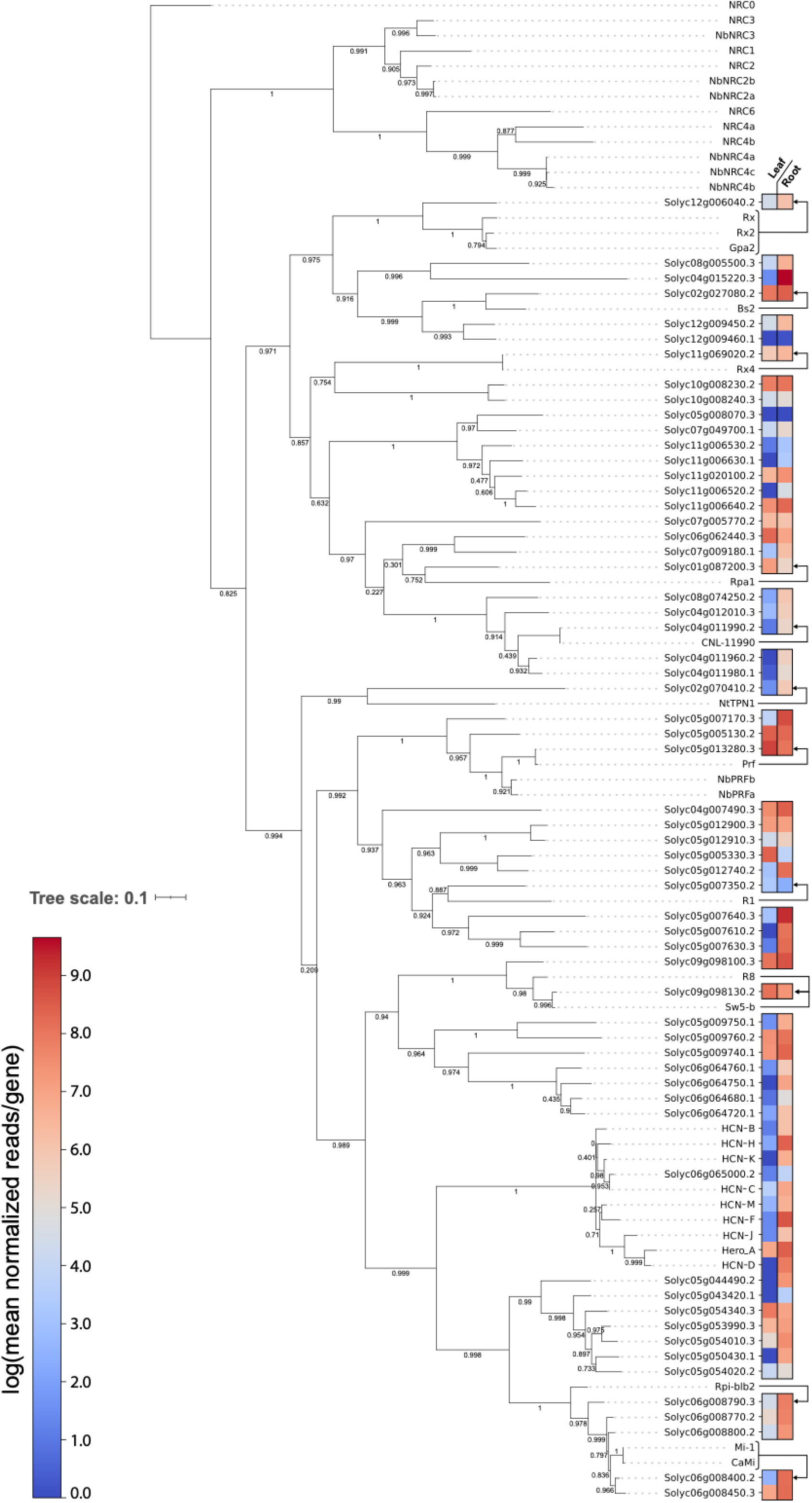
Resistance gene homologs in tomato show varying degrees of expression in roots and leaves. A phylogenetic tree was constructed using selected NRC2/3/4 dependent sensor NLRs from the RefPlant NLR dataset (Kourelis et al., 2021) and the NRC sensor clade of the tomato NLRome. The amino acid sequences of the NB-ARC domain were aligned using MAFFT, and the phylogenetic tree was generated using FastTree. The NRC helper clade served as an outgroup, and the tree was rooted on NRC0. Arrows highlight the closest tomato homolog to the NRC2/3/4 dependent sensor NLRs. The log-values of means for normalized counts, determined by RNA-seq two-week-old unchallenged LA1792 tomato plants, were used to indicate the expression of the respective tomato genes in roots and leaves.

## List of Supplementary data

**Supplementary Data 1.** NLR gene IDs and NLR phylogenetic sub-clades from tomato

**Supplementary Data 2.** NLR gene distance matrix from tomato

**Supplementary Data 3.** Overview and summary of NLR gene clusters from tomato

**Supplementary Data 4.** Genomes used for NRC6 and HCN homolog detection for Figure 2

**Supplementary Data 5.** Log2-fold changes detected in RNA-seq between leaf and root tissues

**Supplementary Data 6.** Raw and normalized counts detected in RNA-seq of leaf and root tissues

**Supplementary Data 7.** Plasmids used and constructed in this study

**Supplementary Dataset 1.** Distance file, alignment and phylogenetic tree files used for Figure 1

**Supplementary Dataset 2.** Sequence, alignment, and phylogenetic tree files used for Figure 2

**Supplementary Dataset 3.** Sequence, alignment, and phylogenetic tree files used for Figure 6

**Supplementary Dataset 4.** HR scoring data used for Figures 3; 4; 5; 6

